# Identification of Subpallial Neuronal Populations Across Zebrafish Larval Stages that Express Molecular Markers for the Striatum

**DOI:** 10.1101/2021.08.11.455880

**Authors:** Vernie Aguda, Helen Chasiotis, Indira Riadi, Tod Thiele

## Abstract

Striatal neurons play a central role in vertebrate action selection; however, their location in larval zebrafish is not well defined. We assayed for conserved striatal markers in the zebrafish subpallium using fluorescent *in situ* hybridization (FISH) and immunohistochemistry. Whole mount FISH revealed an inhibitory neuronal cluster rostral to the anterior commissure that expresses *tac1*, a gene encoding substance P. This molecular profile is shared by mammalian striatal direct pathway neurons. A second partially overlapping population of inhibitory neurons was identified that expresses *penka*, a gene encoding enkephalin. This molecular profile is shared by striatal indirect pathway neurons. Immunostaining for substance P and enkephalin confirmed the presence of these peptides in the subpallium. The *tac1* and *penka* populations were both found to increase linearly across larval stages. Together, these findings support the existence of a striatal homologue in larval zebrafish that grows to match the development and increasing behavioural complexity of the organism.

## 1 Introduction

The basal ganglia (BG) are a collection of brain nuclei located in the ventral or subpallial part of the telencephalon that play a central role in the control of motor behaviour and action selection (Grillner *et al.,* 2013). They are anatomically and neurochemically similar across vertebrate species, indicating that they may perform similar computations and behavioural control across species (Wullimann, 2014). Although primarily known for playing a vital role in motor actions, the BG have also been implicated in addiction, motor learning, and other higher-level cognitive functions. A dysfunction in these nuclei is often associated with movement disorders, such as Parkinson’s Disease (PD) and other dyskinesias and understanding the basis of their clinical manifestations requires detailed knowledge about the function of each neuronal subpopulation within the BG (Wilson, 1925).

### 1.1 Basal Ganglia Anatomy and Basic Circuitry

In humans, the striatum (caudate, putamen, and ventral striatum) and globus pallidus (consisting of internal, GPi, and external, GPe, segments) are telencephalic brain areas that comprise the primary components of the BG. Within these structures there exists dorsal and ventral striatopallidal systems, with the dorsal system associated with motor control and the ventral system involved in limbic functions (Heimer *et al.,* 1995). The subthalamic nucleus (STN) located in the diencephalon and the mesencephalic substantia nigra pars reticulata (SNr) are also classified as components of the BG (Lanciego *et al.,* 2012; Figure 1.1.1). The main input center for the BG is the striatum, which receives afferents from the cortex and thalamus that relay information about body position and environmental circumstances. Afferent inputs influence the activity of GABAergic medium spiny neurons (MSNs) that make up ∼95% of the striatal neuronal population, of which there are two classes. The first class expresses the dopamine type 1 receptor (D1), is excited by dopamine, and projects directly to the output centers of the BG (SNr and GPi). This circuitry represents the ‘direct pathway’ and promotes movement by inhibiting the GPi and SNr, leading to the disinhibition of downstream motor centers. The second class of MSNs expresses dopamine type 2 receptors (D2), is inhibited by dopamine, and sends most of its projections to the GPe. Decreased activity in the GPe in turn leads to greater activity in the STN and GPi, ultimately causing more inhibition on downstream motor areas. (Albin *et al.,* 1989). This circuitry represents the ‘indirect pathway’, and the overall effect of its activity is to inhibit new movements and arrest ongoing movements (Figure 1.1.2).

**Figure 1.1.1:**
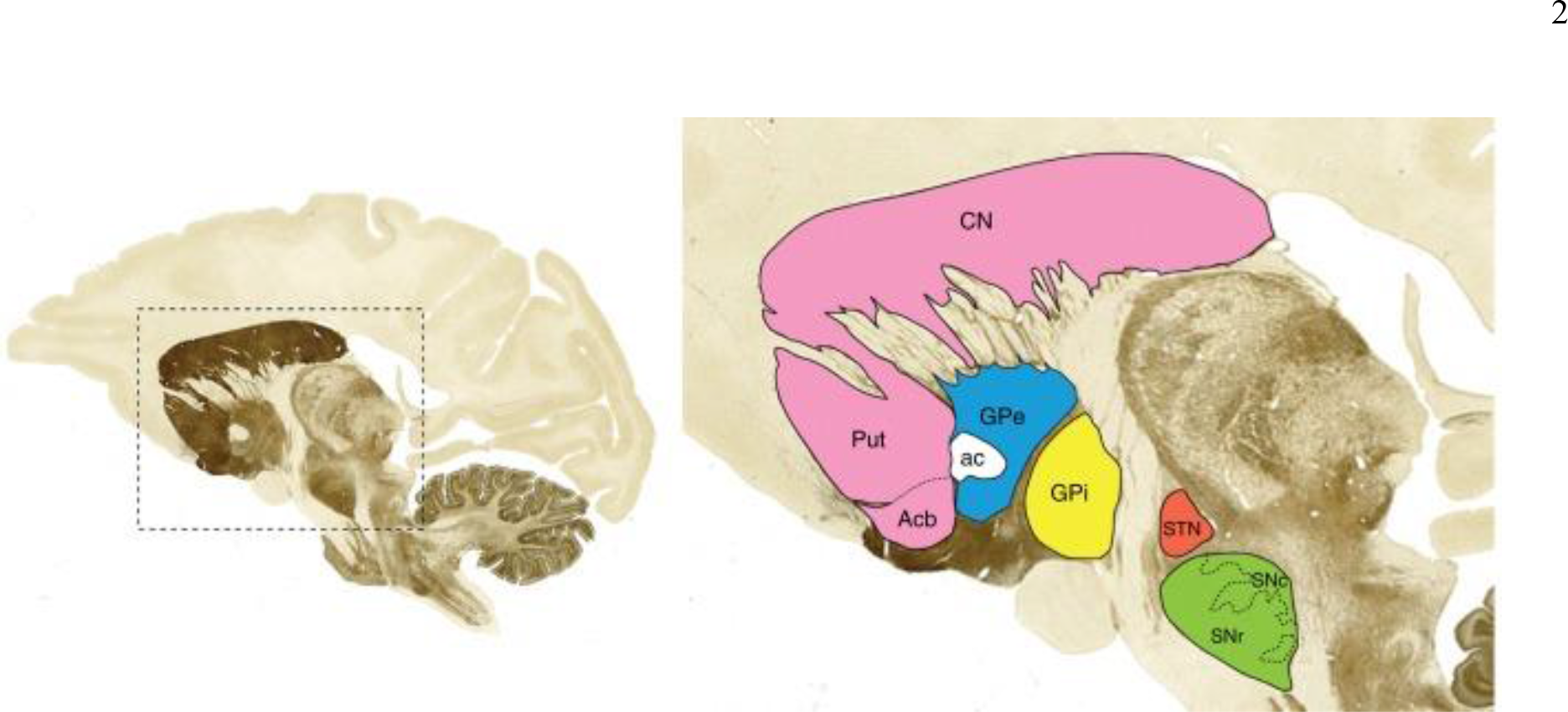
Core mammalian basal ganglia nuclei. Sagittal view of a monkey brain stained with acetylcholinesterase (left) showing the location of the major components of the basal ganglia system (right). Image taken from Lanceigo, Luquin, and Obeso (2012).

**Figure 1.1.2:**
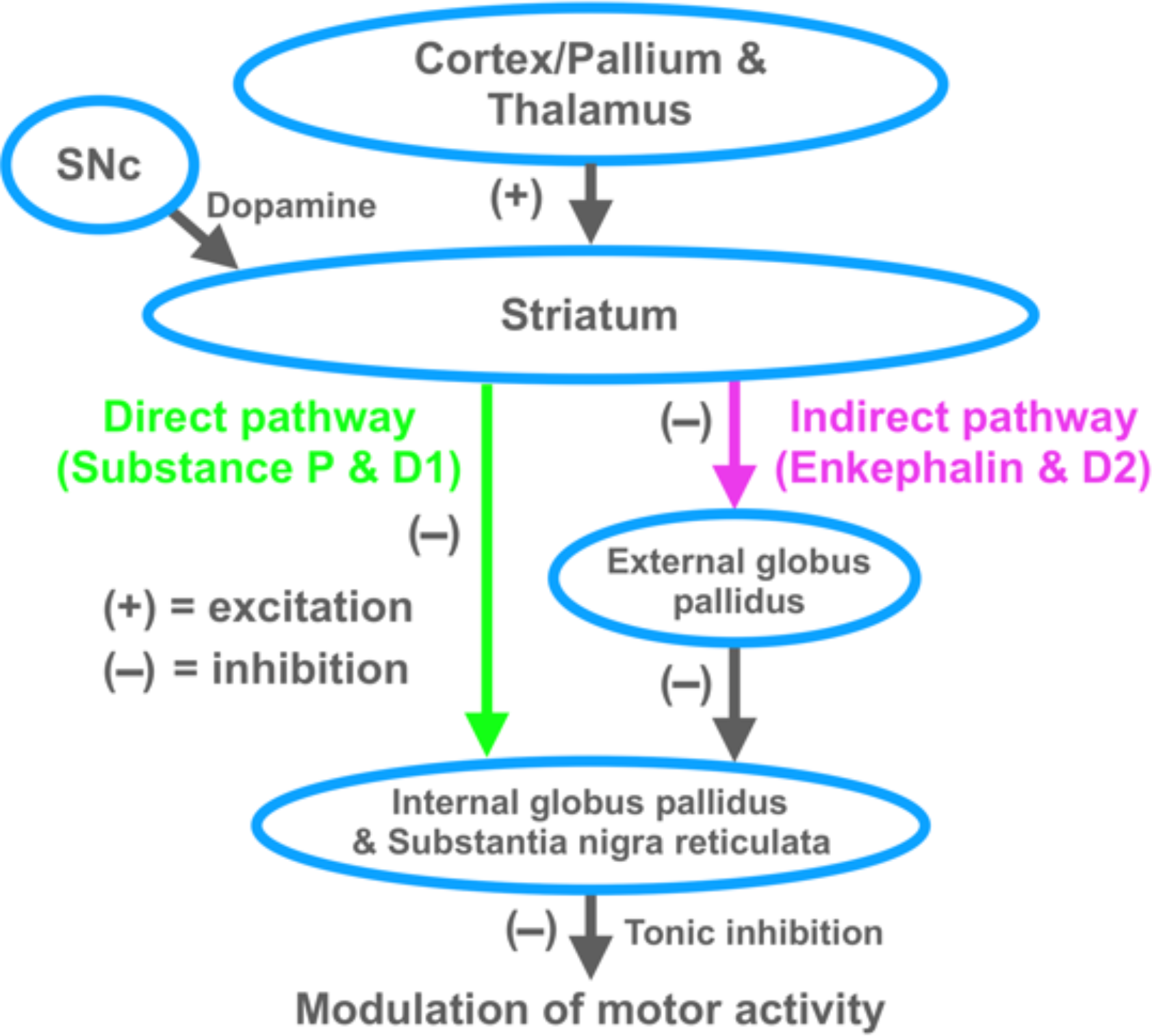
Direct and indirect basal ganglia pathway circuitry. Schematic diagram of basal ganglia circuitry in a mammal highlighting the direct and indirect pathway and the neurotransmitters involved. SNc, substantia nigra pars compacta; D1, dopamine type 1 receptor; D2, dopamine type 2 receptor.

Dopamine (DA) plays a central role in modulating the activity of the striatal output pathways. Mesencephalic dopaminergic neurons within the substantia nigra pars compacta (SNc) provide dopaminergic tone to the two striatal pathways. D1 and D2-expressing MSNs respond to dopamine differently in order to carry out their specific functions of initiating and suppressing movements, respectively. When SNc DA neurons are active, D1-expressing MSNs become further excited, whereas D2-expressing MSNs become inhibited (Stephenson-Jones *et al.,* 2011). This influence on striatal activity is mediated through dopaminergic activation of G-protein-coupled receptors, which either excite (D1) or inhibit (D2) MSNs altering their responsiveness to glutamatergic signaling. D1 receptor stimulation increases the responsiveness of striatal neurons to a sustained synaptic release of glutamate through enhanced opening and upregulation of NMDA receptors (Hallett *et al.,* 2006) and inactivation of K^+^ channels (Snyder *et al.,* 2000). Conversely, D2 receptor stimulation reduces the responsiveness of striatal neurons through decreased AMPA receptor currents (Cepeda *et al.,* 1993), reduced opening of Na^2+^ channels (Surmeier *et al.,* 1992), and enhanced opening of K^+^ channels (Greif *et al.,* 1995).

### 1.2 Basal Ganglia Development and its Genetic Control

Regional specification of the central nervous system (CNS) begins at the neural plate stage during gastrulation. During the development of the telencephalon from the neural tube, the rapid migration of postmitotic neurons in the subpallium form three prominent progenitor domains: the septum, medial ganglionic eminence (MGE), and lateral ganglionic eminence (LGE) (Wilson & Rubenstein, 2000; Corbin *et al.,* 2008; Hebert & Fishell, 2008). The MGE is the most ventrally positioned domain and gives rise to the GP as well as GABAergic neurons that populate the cortex, GP, and striatum. The LGE is more dorsally positioned and gives rise to the caudate and putamen (striatum) (Evans *et al.,* 2012). Most GABAergic MSNs that populate the striatum are also born in the LGE.

Anatomically, the BG can be distinguished by the expression of genetic markers during development. From a developmental standpoint, the striatum and the pallidum represent two subpallial components that originate from the expression of genes involved in the regional specification of the telencephalon. Patterning that occurs along the anterioposterior and dorsoventral axis influences cell fate specification and the generation of distinct neuronal subtypes (Wilson & Rubenstein, 2000). Several homeobox genes are critical for mediating the ventral CNS patterning of the telencephalon and establishing boundaries between regional subdivisions.

In mammals, *nkx2.1* has distinct anterioposterior and dorsoventral boundaries and is required for the development of pallidal-related structures (Sussel *et al.,* 1990). In KO mouse studies, there is a molecular repatterning of the MGE into LGE-like tissue, leading to an enlargement of LGE derivatives later in gestation. This also causes a reduction of interneurons and GABAergic neurons in the cerebral cortex (Sussel *et al.,* 1990). In non-vertebrates, fly embryos lacking *vnd* (ventral nervous system defective, which encodes for the NK2 homeodomain protein) show a ventral-to-dorsal transformation, mirroring the phenotype of *nkx2.1* mammalian mutants (McDonald *et al*., 1998; Weiss *et al.,* 1998). Counter expression of *gsh2* and *pax6* form the border between the dorsal and lateral region of the telencephalon and reciprocal regulatory interactions between these genes mediate dorsoventral patterning of the pallial/ subpallial boundary (Corbin *et al.,* 2003). Mutant *gsh2* mice contain pallial markers in the dorsal LGE, indicating dorsalization, whereas mutant *pax6* mice show subpallial identity within the ventral pallium. The specification of the striatum depends on the function of *gsh1/2*, which drives expression of *dlx1/2* and *mash1*. These two genes work in conjunction, each repressing the expression of the other to drive LGE development and identity (Long *et al.,* 2009). Mutant *dlx1/2* mice express ventral pallial and MGE transcription factors in the LGE, have lower levels of *GAD67*, and have severe defects in striatal and olfactory tubercle development (Long *et al.,* 2007). This suggests that *dlx* genes are essential for repressing dorsal and ventral identities in the LGE and promoting GABAergic cell fate. Several rodent studies have also revealed that the specific birth date of a neuron defines the segregation of striatal neurons of the striosomes and the matrix, which are involved in anatomical and functional compartmentalization within the striatum (van der Kooy & Fishell, 1987; Krushel *et al.,* 1993). This suggests that neuronal cell fate within the BG is determined by birth order and lineage relationships dictated by genes that control forebrain developmental patterning.

### 1.3 Conservation of Basal Ganglia Structures Across Vertebrate Lineages

The core structures of the BG are highly conserved across vertebrate evolution. There are multiple studies providing evidence that the striatum and pallidum were present in early vertebrate lineages (Parent *et al*., 1986; Smeets *et al.,* 2000). The mammalian striatum is often distinguished by an abundance of GABAergic MSNs (Parent *et al.,* 1986; Medina & Reiner, 1995). These neurons contain either substance P (SP) or enkephalin (ENK) and project to the pallidum. Other characteristics of the striatum include: an abundance of dopaminergic and cholinergic terminals, distinct types of interneurons (cholinergic interneurons, GABAergic interneurons containing calretinin, parvalbumin, and nitric oxide synthase), low firing rates, and the presence of DA neurons for rodents and primates (Graybiel, 1990; Marin *et al.,* 2005). The pallidum is characterized by GABAergic projection neurons containing LANT6, SP and ENK wooly fibres, and scarce dopaminergic fibres. (Harber & Nauta, 1983; Graybiel, 1990)

#### 1.3.1 Anamniotes

In birds, reptiles, amphibians, and some fish, the BG shows the same basic neurochemistry, connectivity, and genetic embryological patterns as in mammals, supporting the idea that the BG is a common vertebrate structure for the control of movement (Figure 1.3.1; Russchen *et al.,* 1987; Medina & Reiner, 1995). In the ancient jawless lamprey, the oldest lineage of vertebrates that diverged 560 million years ago, a ventromedial zone of the telencephalon, akin to the subpallial region of most jawed vertebrates, contains GABAergic neurons that express D1 receptors and are enriched with ENK (Pombal *et al.,* 1997) and GABAergic neurons that express D2 receptors and are enriched with SP (Auclair *et al.,* 2004). Moreover, this region also expresses lamprey *dlx1/2*, and receives DA input from the posterior tubercle of the diencephalon and midbrain (Myojin *et al.,* 2001; Pierre *et al.,* 1994).

**Figure 1.3.1:**
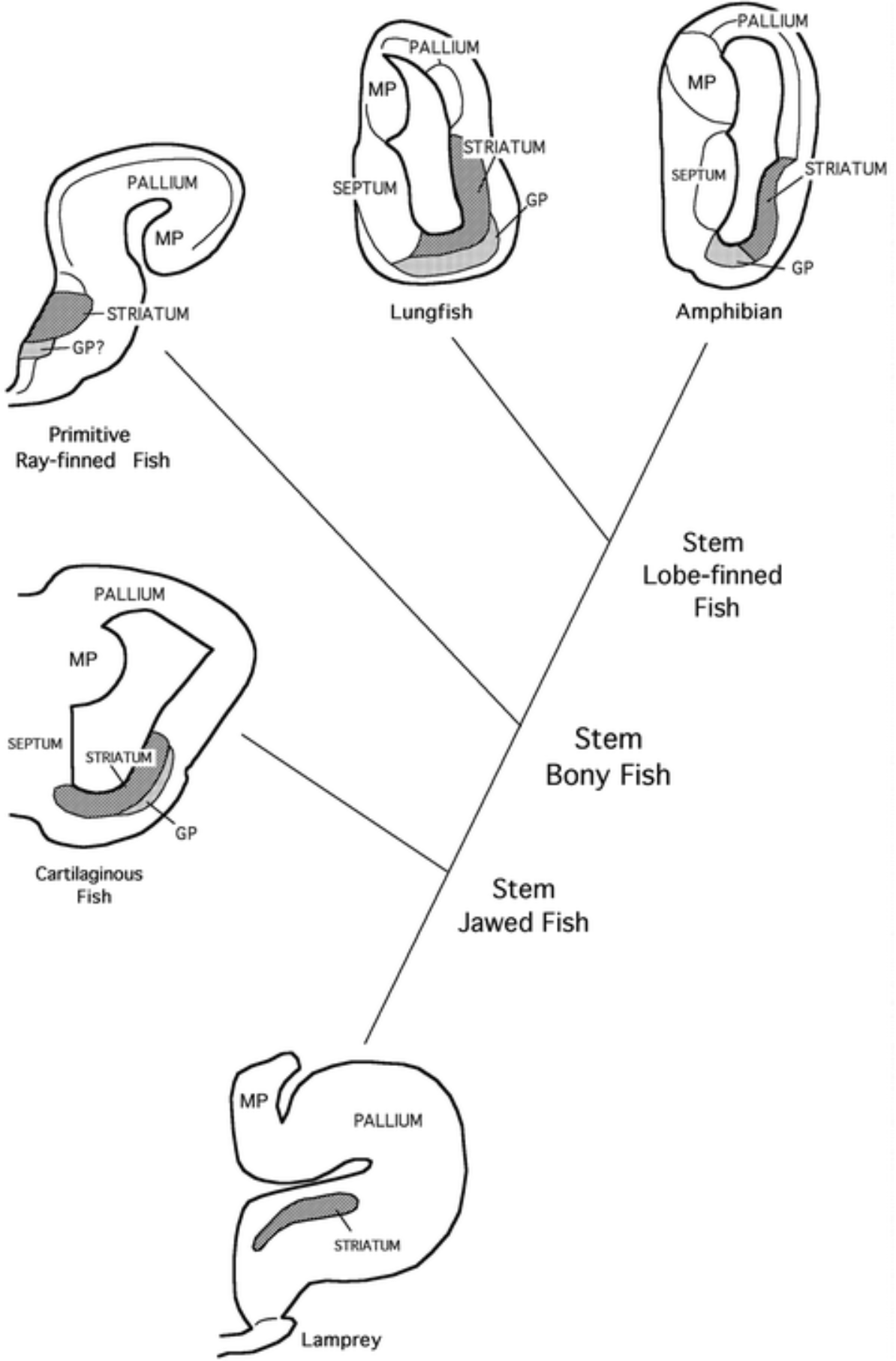
The position of the striatum and pallidum across anamniotes. Schematics of frontal sections through the BG of five anamniote groups. Note that the BG in all groups with an evaginated telencephalon have their striatum and GP located around a lateral ventricle. In ray-finned fish with an everted telencephalon, the striatum is thought to be located closer to the midline. Image adapted from Reiner (2009).

Cartilaginous fish (chondroichthyes), among the most ancient living representatives of the jawed vertebrates, also contain striatal and pallidal progenitor domains within their ventral telencephalon (Hedges, 2001). The striatal domain, corresponding to the LGE in mammals, expresses *dlx2* but not *nkx2.1*, whereas both *dlx2* and *nkx2.1* are expressed in the pallidal domain, which corresponds to the MGE in mammals (Medina *et al.,* 2014). Bony fish (osteicthyes) diverged into ray-finned fish (actinopterygians) and lobe-finned fish (sarcopterygians), with one of the hallmark differences being the pallial eversion of ray-finned fish during development (Northcutt & Braford, 1980). This eversion reverses the medial to lateral topography of the pallium but the topographical relations among telencephalic cell groups remain similar. Gene expression analysis, neurochemical characterization of SP and ENK neurons, and the presence of dopaminergic innervation indicate that the dorsal region of the ventral area of the subpallium (Vd) may be the striatal homolog in ray-fined fish (Reiner & Northcutt, 1992). The location of a pallidal homolog is believed to either reside as an intermingled population among the striatal neurons of the Vd, as evidenced through *nkx2.1* expression, or found in a ventral part of the ventral area (Vv), which also expresses *nkx2.1* and is in the same relative topographical position of the mammalian MGE (Alunni *et al.,* 2004). In lobe-finned fish, the presence of a striatum and GP in the ventrolateral telencephalon has been characterized by the presence of GABAergic MSNs, SP and ENK neurons, dopaminergic innervation from midbrain structures, and region-specific expression of *dlx2* and *nkx2.1* in the striatum and pallidum, respectively (Reiner & Northcutt, 1987, Vallarino *et al.,* 1998). The ventrolateral telencephalon of amphibians also contains a striatal sector, containing SP and ENK neurons, GABAergic neurons (Inagaki *et al.,* 1981; Taban & Cathieni, 1983), and *dlx1/2* expression (Papalopulu & Kintner, 1993). The pallidal area, found in the caudal ventrolateral telencephalon, contains a large population of GABAergic neurons and expresses *nkx2.1* (Marin *et al.,* 1997). Dopaminergic innervation of the striatal sector arises from the posterior tubercle and the midbrain (Dube & Parent, 1990; Gonzalez & Smeets, 1991).

#### 1.3.2 Amniotes

Through the course of evolution, the expansion of both the pallium and subpallium caused several changes to occur in the BG organization to accommodate for the increasing number of telencephalic neurons. Tegmental DA input into the striatum became more substantial and motor control occurred via pallial motor areas rather than directly to the pretectal region (Reiner, 2016). In mammals, the evolution from a simple pallium to a multilayered neocortex led to the BG occupying a more central position within the telencephalon due to extensive lateroventral migration from pallial neurons (Karten, 1969). By contrast, in reptiles and birds the striatum is ventrally located with the pallial regions just dorsal to it. Histochemical studies show that the ventrolateral telencephalon of reptiles contains both a striatum, as evidenced by SP and ENK GABAergic neurons (Reiner, 1987), expression of *dlx1/2* (Smith-Fernandez *et al.,* 1998), and a globus pallidus, showing *nkx2.1* expression. The overlap of SP and ENK wooly fibres in the globus pallidus suggest that GPi and GPe neurons are intermingled in the reptile (Reiner, 1987). Further telencephalic enlargement is seen in the avian telencephalon, which shows distinct compartmentalization of some BG components that is not present in reptiles (Reiner, 1993).

Striatal neurons expressing SP showed to preferentially project to the GPi, whereas striatal neurons expressing ENK preferentially project to the GPe (Anderson & Reiner, 1990). The presence of SP and ENK GABAergic neurons, and *dlx1/2* classify the striatal region (Smith-Fernandez *et al.,* 1998), and the globus pallidus is identifiable by *nkx2.1* expression, and GABAergic neurons containing LANT6 and PARV (Karten & Dubbeldam, 1973).

In mammals, the striatum is compartmentalized into zones called striosomes, which are embedded in the matrix (Graybiel, 1990). Neurons populating these two sectors differ in connectivity and neurochemical make-up, with striosomal neurons preferentially expressing SP and connecting to the GPi and tegmental dopaminergic neurons while neurons in the matrix preferentially express ENK and connect to the GPe (Charron *et al.,* 1995). The neurons in the striosome are poorer in acetylcholinesterase (AChE), richer in mu-opiate receptors, and poorer in calbindin than the neurons in the matrix (Graybiel & Ragsdale, 1978). The presence of SP and ENK GABAergic neurons, *dlx* expression, and dopaminergic input also identify the striatal component of the BG in mammals (Graybiel *et al.,* 1981; Gerfen, 1992). The GP is characterized by *nkx2* expression, GABAergic neurons containing LANT6 and PARV, and the presence of SP and ENK wooly fibres (Haber & Nauta, 1983). By recognizing the conserved traits and evolutionary changes in telencephalic morphology, connectivity, and neurochemistry, salient conclusions about brain regions and their underlying circuitry across species can be made (Figure 1.3.2).

**Figure 1.3.2:**
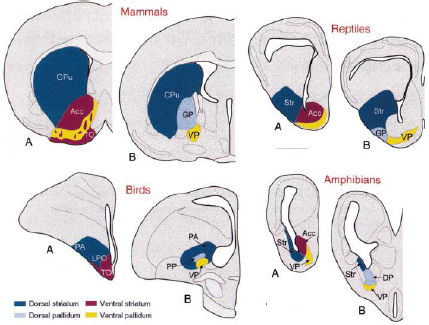
The position of the striatum and pallidum across amniotes. A schematic of transverse sections at rostral (A) and caudal (B) telencephalic levels to illustrate the relative position of striatal and pallidal structures across tetrapods. Acc, nucleus accumbens; Cpu, caudate-putamen; DP, dorsal pallidum; GP, globus pallidus; LPO, lobus parolfactorius; Str, striatum; VP, ventral pallidum. Image adapted from Smeets (2000).

### 1.4 Functional Studies of Basal Ganglia Circuitry

Historically, BG research has been guided by a model of the direct pathway projections from the striatum promoting movement and indirect pathway projections inhibiting movement (Albin *et al.,* 1989; Delong, 1990). This model, guided by what was known about BG anatomy at the time, was revised to account for the presence of the striosomes and matrix (Graybiel *et al.,* 1981). In mammals, it is understood that there are subpopulations of striatal projection neurons that have different projection targets and dopamine effects these subpopulations differently. Direct pathway neurons are characterized by expressing the D1 type dopamine receptor and containing the neuropeptides substance P and dynorphin. Indirect pathway neurons are characterized by expressing the D2 type dopamine receptor and containing the neuropeptide enkephalin (Gerfen *et al.,* 1990; Le Moine *et al.,* 1995). Direct pathway neurons are shown to innervate mainly the SNr and GPi, whereas indirect pathway neurons primarily innervate the GPe (Albin *et al.,* 1989).

Advancements in technology and research techniques have expanded the study of brain anatomy, connectivity, and circuitry. In mammals, the direct and indirect pathways have been suggested to control limbic functions in addition to motor selection, and recent work involving rodents and optogenetics have allowed for specific clusters of BG neurons to be activated or silenced in order to determine their role in action selection. Optogenetic experiments have validated the classical BG model of D1-MSN activation leading to movement and D2-MSN activation leading to cessation of movement (Kravitz *et al.,* 2010). However, additional studies provide evidence for activity in both D1-MSNs and D2-MSNs during movement, and low activity in both populations during immobility (Cui *et al.,* 2013; Isomura *et al.,* 2013). Other work has provided further evidence that the two pathways work together to facilitate desired movements while simultaneously suppressing competing ones (Mink, 1996; Hikosaka *et al.,* 2000). Despite all this recent progress, there still lacks a comprehensive analysis on these MSNs due to the dense neural network in which MSNs are embedded (Gerfen, 1992).

Until recently, it was unknown whether similar BG circuitry existed between mammals and basal vertebrates. A study done by Stephenson-Jones (2011) and colleagues showed not only the presence of mammalian BG components in lamprey, but also that circuit features and physiological activity patterns were conserved. The direct and indirect striatopallidal pathways, nigrostriatal pathways, as well as GP projections to the thalamus and downstream locomotor regions were all shown to be conserved. Most BG output in the lamprey is focused on brainstem motor areas, suggesting that the primary function of the BG is to modulate action selection via these motor areas. A similar situation appears in birds, where dopaminergic input to the striatum provided by the SNc helps promote movement, as seen by administration of DA agonists producing stereotypic pecking or vocalization (Akbas *et al.,* 1984).

### 1.5 Larval Zebrafish as a Model for Basal Ganglia Development and Function

Larval zebrafish offer several advantages for examining the structure and function of neural circuits. Their brain has a medium complexity and can be functionally imaged and modulated with cellular resolution across all brain regions. They are relatively easy to maintain, cost efficient, have quick generation periods, and remain transparent at early life stages allowing for unobstructed imaging in behaving animals without surgery. Powerful genetic tools, such as the Gal4/UAS system and CRISPR/ Cas9 genome editing are also well established in this model. Zebrafish engage in stereotypical behaviours at the larval stage, such as the optomotor response, prey capture, and predator avoidance (Budick & O’Malley, 2000). This coupled with functional imaging techniques allows for causal relationships to be made between neuronal activity and behavior production. Due to the highly conserved vertebrate brain plan throughout evolution, zebrafish larvae contain many of the same brain structures as higher vertebrates (Wullimann, 2014).

A significant amount of work has gone into analyzing gene expression patterns of canonical striatal and pallidal markers such as *dlx1/2*, *nkx2.1*, and *GAD67* in the teleost class of fish to which zebrafish belong (Ganz *et al.,* 2012). There are parallels in the distribution of cholinergic cells (Parker *et al.,* 2013), presence of dopaminergic D1 and D2 receptors (Le Crom *et al.,* 2003), and ascending dopaminergic input (Rink & Wullimann, 2001) between mammals and zebrafish. Whether a mammalian striatal homolog is present and functional in larval zebrafish remains an open question. Identifying a BG homolog is potentially complicated by the partially everted nature of the teleost forebrain. All other vertebrate telencephalons undergo invagination during development (Folgueira *et al.,* 2012). Eversion arises through a lateral out folding of the medial and dorsal pallium, placing the hippocampus and olfactory cortex at a more lateral position than one would find in mammals (Figure 1.5.1). Although the effects of eversion are strongest in the pallium, there is potential for some effect on the subpallium as well. In most vertebrates, striatal and pallidal structures are located near the lateral ventricles, but in teleosts the location of these structures is thought to lie more medially (Reiner, 2009).

**Figure 1.5.1:**
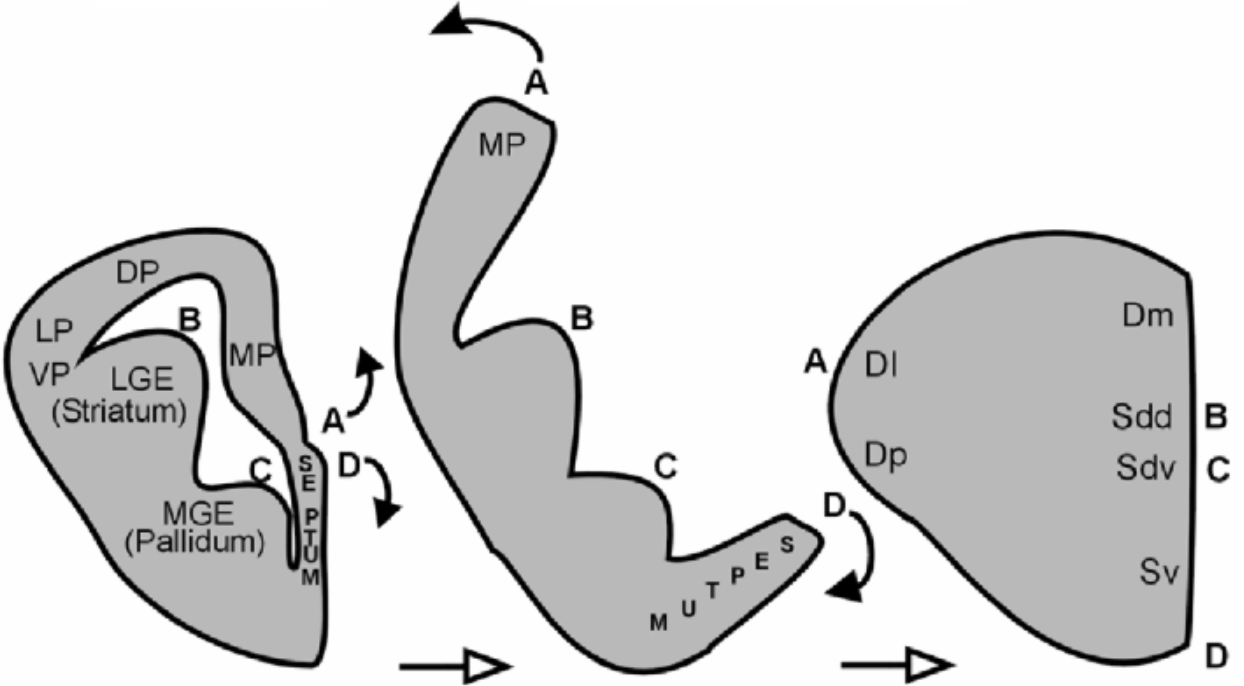
Theoretical mapping of an early mouse telencephalon onto the zebrafish telencephalon. Schematic illustrating the partial eversion of the medial pallium and dorsal pallium. Left: transverse view of an early mouse telencephalon. Middle: depiction of the eversion process. Right: transverse view of the zebrafish telencephalon DP, dorsal pallium (isocortex); LGE, lateral ganglionic eminence; LP, lateral pallidum (olfactory cortex); MGE, medial ganglionic eminence; MP, medial pallium (hippocampus); Sdd, dorsal subdivision of dorsal subpallium (“striatum”); Sdv, ventral subdivision of dorsal subpallium (“pallidum”); Sv, ventral subpallium (septum); VP, ventral pallidum. Image adapted from Wullimann (2009).

### 1.6 Thesis Goals and Aims

#### 1.6.1 Goal

We intend to determine if larval zebrafish contain circuitry homologous to the mammalian striatum through a striatal gene expression analysis of neurons within the subpallium. We will examine time points that cover the development of behavior from purely reflexive behaviour to a stage when complex behaviours such as sociality exist. This gene expression analysis will provide important insights into striatal development and subpallial compartmentalization within a model system that has unique advantages for understanding brain function.

## Methods

### 2.1 Zebrafish Maintenance

All adult zebrafish were housed in a facility room held under a 14hr/10hr light/ dark cycle at 28°C and were fed twice a day (Gemma micro 300 ZF). Adult crosses included 2-3 male and 1-2 female fish. Larvae were collected the day of fertilization, placed in Danieau’s solution and kept in an incubator held at 28°C. From 5 days post fertilization (dpf) onward, larvae were fed Gemma micro 75 ZF. Fish past 7 dpf were transferred to a 3L tank and placed on the system. For all experimental procedures, larvae were sacrificed by placing them in ice cold water prior to tissue collection and fixation. All animal experiments were performed with the approval of the University of Toronto Animal Care Committee in accordance with the guidelines from the Canadian Council for Animal Care (CACC).

### 2.2 Molecular Techniques

#### 2.2.1 Fixation and storage

Samples were fixed overnight in 4% paraformaldehyde (PFA) (Electron Microscopy Sciences, 15700) in 0.1M phosphate buffered saline (PBS) (Sigma-Aldrich, P5244) and 4% sucrose (Bioshop, SUC507) at 4^0^C. Brains were dissected manually using forceps and samples were transferred to a scintillation vial with 100% methanol and stored at -20^0^C until further processing.

#### 2.2.2 *In situ* hybridization

Probe generation and whole mount fluorescent *in situ* hybridization (*in situ*) were performed as previously described (Thisse & Thisse, 2014). Briefly, 3, 7, 14, or 21 dpf wild type larvae or similarly aged extracted brains were rehydrated, washed with 1x PBST (PBS with 0.1% Tween-20 (TWN510.500, BioShop)), and their heads were separated. The heads or brains were then digested with 10ug/mL Proteinase K (Roche, RPROTK-RO) for 10-20 minutes at room temperature, post-fixed with 4% PFA in PBST for 20 minutes and washed again with 1x PBST. Samples were incubated at 58^0^C for 2 hours in hybridization mix (50% deionized formamide, SSC(5x), 0.1% Tween 20, 50μg/mL heparin, 500μg/mL RNase-free tRNA, pH 6), and left to incubate for 1-2 nights with 0.25-1ng/μL RNA probe in hybridization mix. The samples were washed with 50% formamide (11814320001, Sigma-Aldrich) in 2X SSCT, 2X SSCT, 0.2X SSCT, and 1x TNT (10mM Tris-HCl, pH 8.0, 150mM NaCl, 0.05% Tween-20). Samples were incubated in blocking buffer (2% blocking reagent (Roche, 11096176001) in 1x TNT) for 2 hours at room temperature and left to incubate for 24 hours with blocking buffer containing a 1:200 dilution of anti-DIG (Roche, 11207733910) or anti-fluorescein antibody conjugated with POD (Roche, 11426346910) at 4^0^C. Next, the samples were washed with 1x TNT and incubated with Cy3 TSA reagent (NEL744001KT, Perkin Elmer) or fluorescein TSA reagent (NEL701A001KT, Perkin Elmer) for 45-60 minutes at room temperature. Samples were washed with 1xPBST then counterstained with DAPI (Sigma-Alrich, D9542, 30ng/μL in PBST) for 2 hours at room temperature. Larvae were finally washed with 1xPBST, mounted in 2% low-melting point agarose (ThermoFisher, 16520050, diluted in 1x Danieu’s solution), and imaged using a 20X 1.0NA water immersion objective on our Zeiss LSM800 confocal microscope. All images were taken through the Zen computer software and viewed using FIJI (Schindelin, 2012). Primers that were used to create RNA probes are summarized in Table 2.2.1.

**Table 2.2.1:**
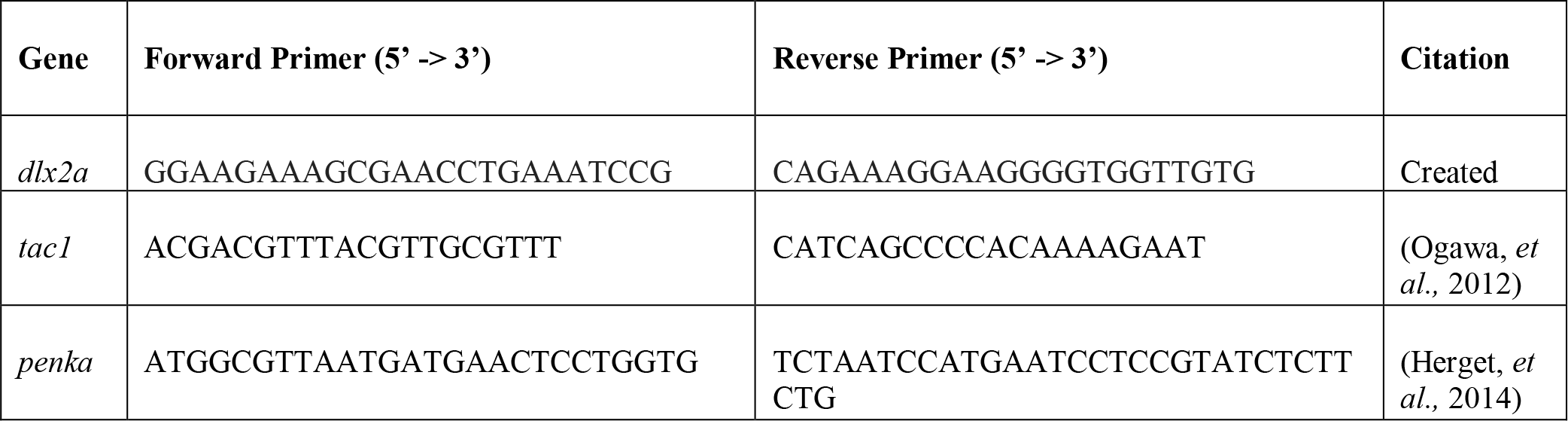
Primers used to create *in situ* probes.

#### 2.2.3 Immunofluorescence

Immunofluorescence (IF) experiments were performed as described previously (Xiao & Baier, 2007). Briefly, larval heads or brains were incubated with primary antibodies for 2 days at 4^0^C, washed with 1x PBST, and incubated with secondary antibodies and DAPI (Sigma-Alrich, D9542, 30ng/uL in PBST) for 2 hours at room temperature. Samples were then washed with 1x PBST and mounted in 2% low-melting agarose (ThermoFisher, 16520050, diluted in 1x Danieu’s solution). The primary antibodies that were used were: rabbit anti-RFP (Rockland, 600-401-379S, diluted to 1:300), rabbit anti-Substance P conjugated Alexafluor 488 (Santa Cruz, sc-21715, diluted to 1:100), rabbit anti-TH (Sigma-Aldrich, AB152, diluted to 1:300), mouse anti-enkephalin (Abcam, ab150346, diluted to 1:50), and mouse anti-ERK (p44/42 MAPK (Erk1/2) (Cell Signalling, L34F12, diluted to 1:500)). The secondary antibodies that were used were: goat anti-rabbit Alexafluor 488 (Invitrogen, A-11034, diluted to 1:300), goat anti-rabbit Alexafluor 594 (Invitrogen, A-11032, diluted to 1:300), goat anti-mouse Alexafluor 488 (Invitrogen, A11029, diluted to 1:300), and goat anti-mouse Alexafluor 598 (Invitrogen, A11032, diluted to 1:300).

### 2.3 Data Analysis

#### 2.3.1 Image Processing, Cell Counting, and Statistics

All images were viewed and processed using FIJI (Schindelin *et al.,* 2012). Each z-stack and z-projection was optimized using image editing techniques included in the program, such as brightness and contrast alterations and the removal of unnecessary background noise. All images were processed in a similar fashion. Cell counts were done manually using the FIJI cell counter tool. Student’s two-tailed unpaired t-tests and regression analysis were performed between different ages of samples within *tac1* and *penka* groups and between *tac1* and *penka* at the same timepoints. The use of multiple comparisons was corrected for using the Bonferroni correction.

### 2.4 Gal4 Driver Line Creation

Putative dopaminergic D1 and D2 receptor lines were created using CRISPR methods described previously (Kimura *et al.,* 2016; Figure 2.4.1). The D1 line was created by the Zebrafish CORE Facility, located within SickKids (Figure 2.4.2). For the D2 line, we constructed guide RNA (gRNA) using ChopChop (http://chopchop.cbu.uib.no/) which cuts the *drd2a* gene ∼600bp upstream of the start codon (gRNA 5; Figure 2.4.3). To summarize, gRNA 5 (250ng) was co-injected with a plasmid containing the donor DNA (Gbait-hsp70:Gal4), gRNA to linearize the plasmid (SgG), and Cas9 RNA into *UAS:Kaede* embryos at the one cell stage. Embryos were placed in an incubator maintained at 26^0^C and screened at 3 dpf for green fluorescence. Larvae that express Kaede in the forebrain were raised on the system and out-crossed to identify potential founders.

**Figure 2.4.1:**
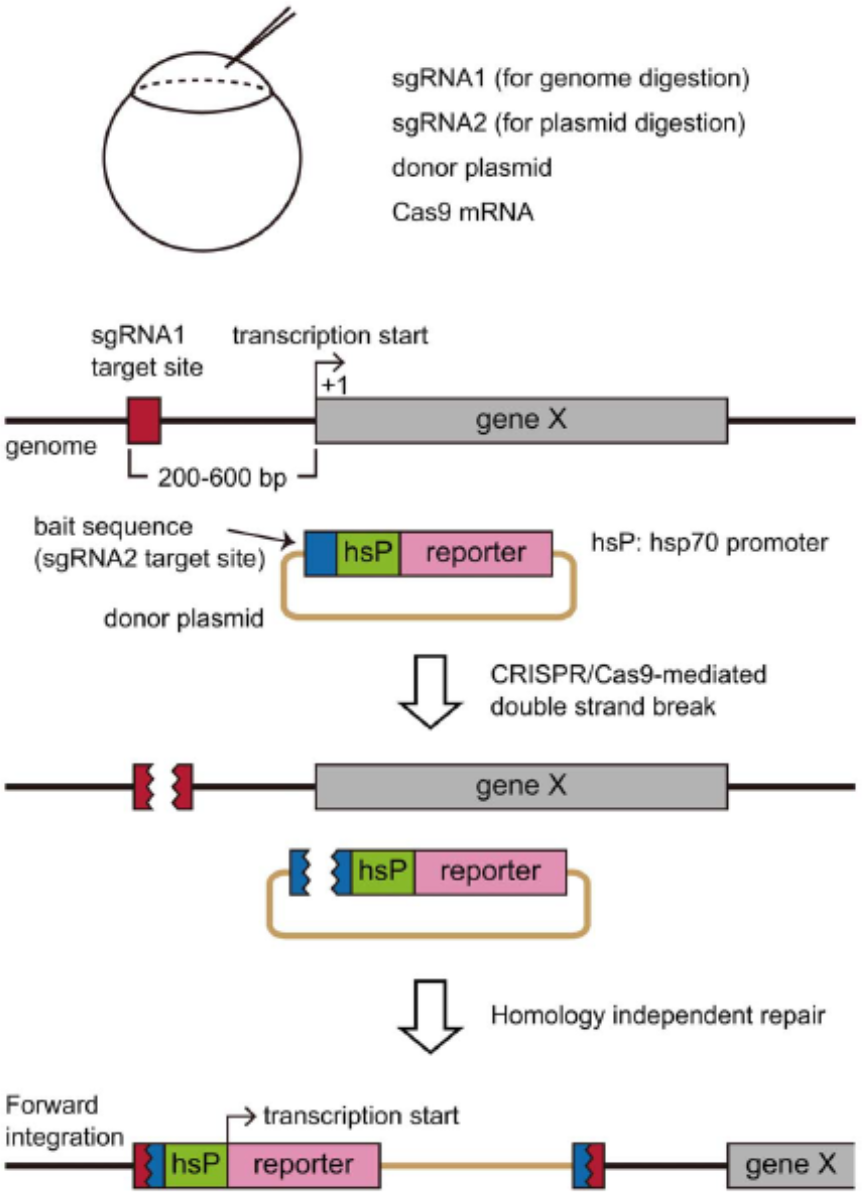
Schematic of CRISPR/Cas9 genome editing. Co-injection of sgRNA1 (for genome digestion), sgRNA2 (for plasmid digestion), donor plasmid, and cas9mRNA into one-cell zebrafish embryos occur. Cas9 mediated cleavage occurs upstream of the gene of interest (∼200-600bp, gene X), as well as the bait sequence on the donor plasmid. This results in the integration of the donor plasmid into the target locus. Adapted from Kimura, *et al.,* (2014).

**Figure 2.4.2:**
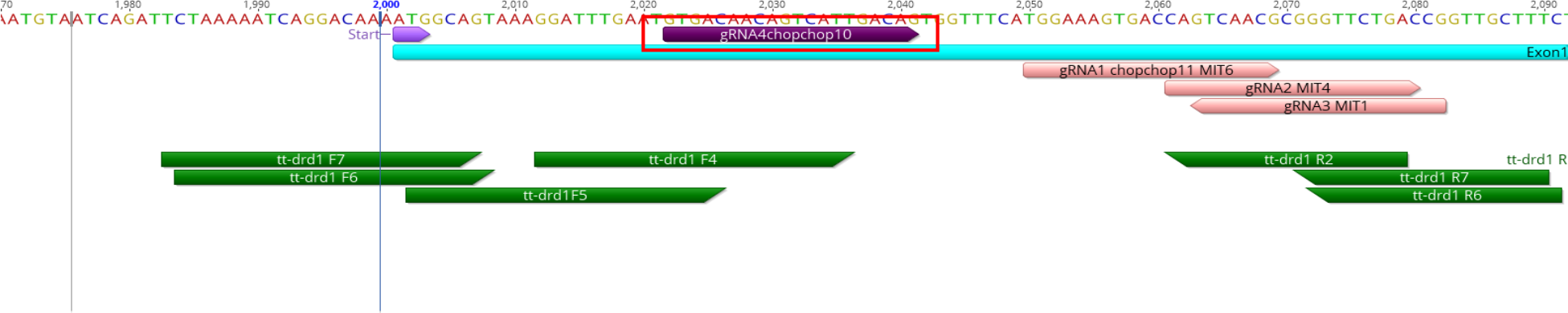
Genome map of the position of guide 4. CRISPR gRNA4 used to make the unverified *drd1a:Gal4;UAS:Kaede* line denoted by red square. Pink arrows represent additional gRNA locations. Turquoise rectangle represents exon 1 of *drd1a*. Green arrows represent PCR primers to validate insertion of gRNA4 into the genome.

**Figure 2.4.3:**
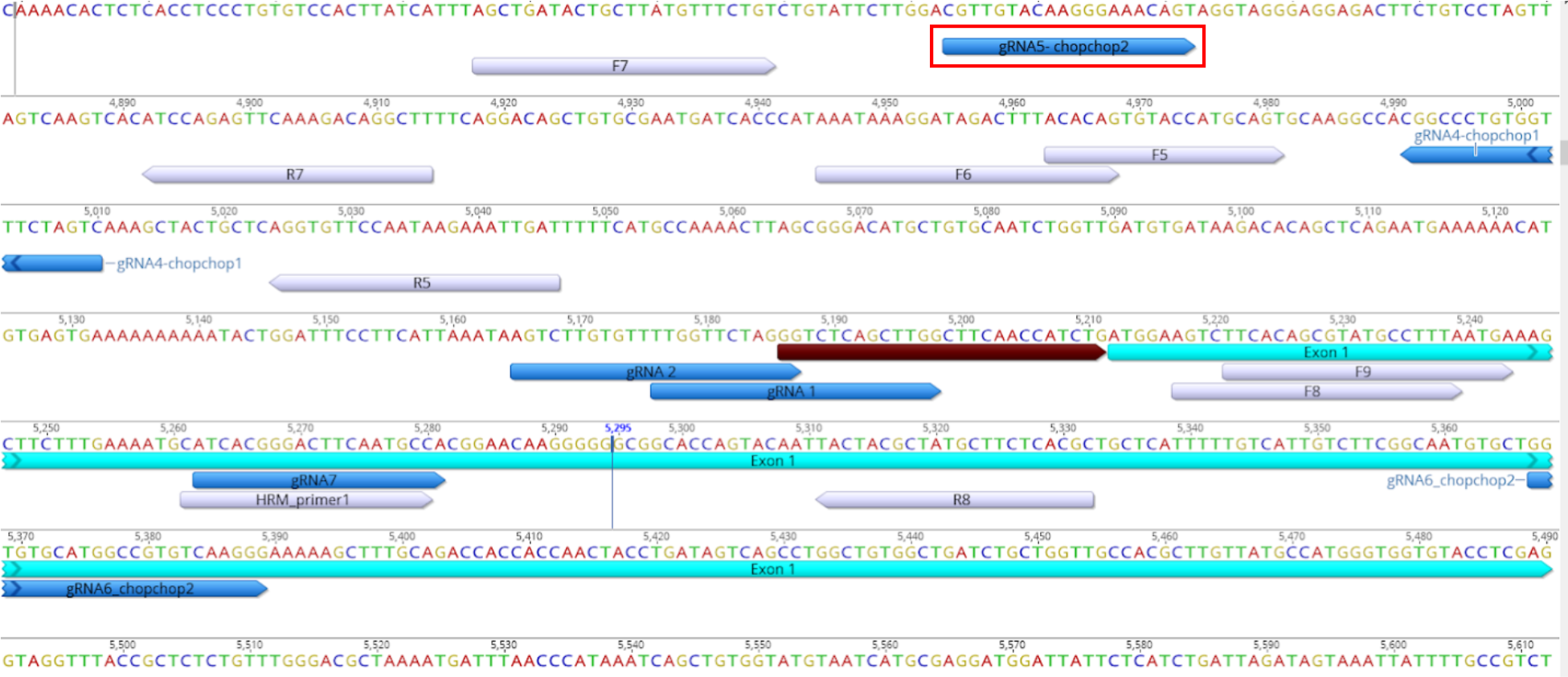
Genome map of the position of guide 5. CRISPR gRNA5 used to make unverified *drd2a:Gal4;UAS:Kaede* line denoted by red square. Brown arrow represents the untranslated region (UTR). Turquoise rectangle represents exon 1 of *drd2a*. Blue arrows represent additional gRNA locations. Light blue arrows represent PCR primers to validate insertion of gRNA into the genome.

### 2.5 Brain Registration

We are currently in the process of fine tuning the digital registration of brains across experiments. For this method, a reference stain of either DAPI (Sigma-Alrich, D9542) or MAPK (Cell Signalling, L34F12) immunostaining is used to morph brains to match morphology. We are using Advanced Normalization Tools (ANTs) for the creation of our brain registry (Marquart, 2017).

## Results

### 3.1 Location of the Subpallium in Larval Zebrafish

The striatum is a subpallial structure, whose GABAergic neurons primarily originate from the LGE during development (Evans *et al.,* 2012). Whole mount *dlx2a* fluorescent *in situ* hybridization was used to validate the location of the larval zebrafish subpallium at 3 dpf (n = 2). In animals probed for *dlx2a* expression, we observed strong expression in the dorsal and ventral subpallium (Figure 3.1.1, B) that extended caudally to the anterior commissure (Figure 3.1.2). Ventral and caudal to the anterior commissure, there was staining of the preoptic region and the ventral thalamus, identifiable by the discrete wing-shaped expression (Figure 3.1.2, D). The resulting expression pattern is consistent with published *dlx2a* staining at 2 dpf performed on brain sections (Mueller, 2008).

**Figure 3.1.1:**
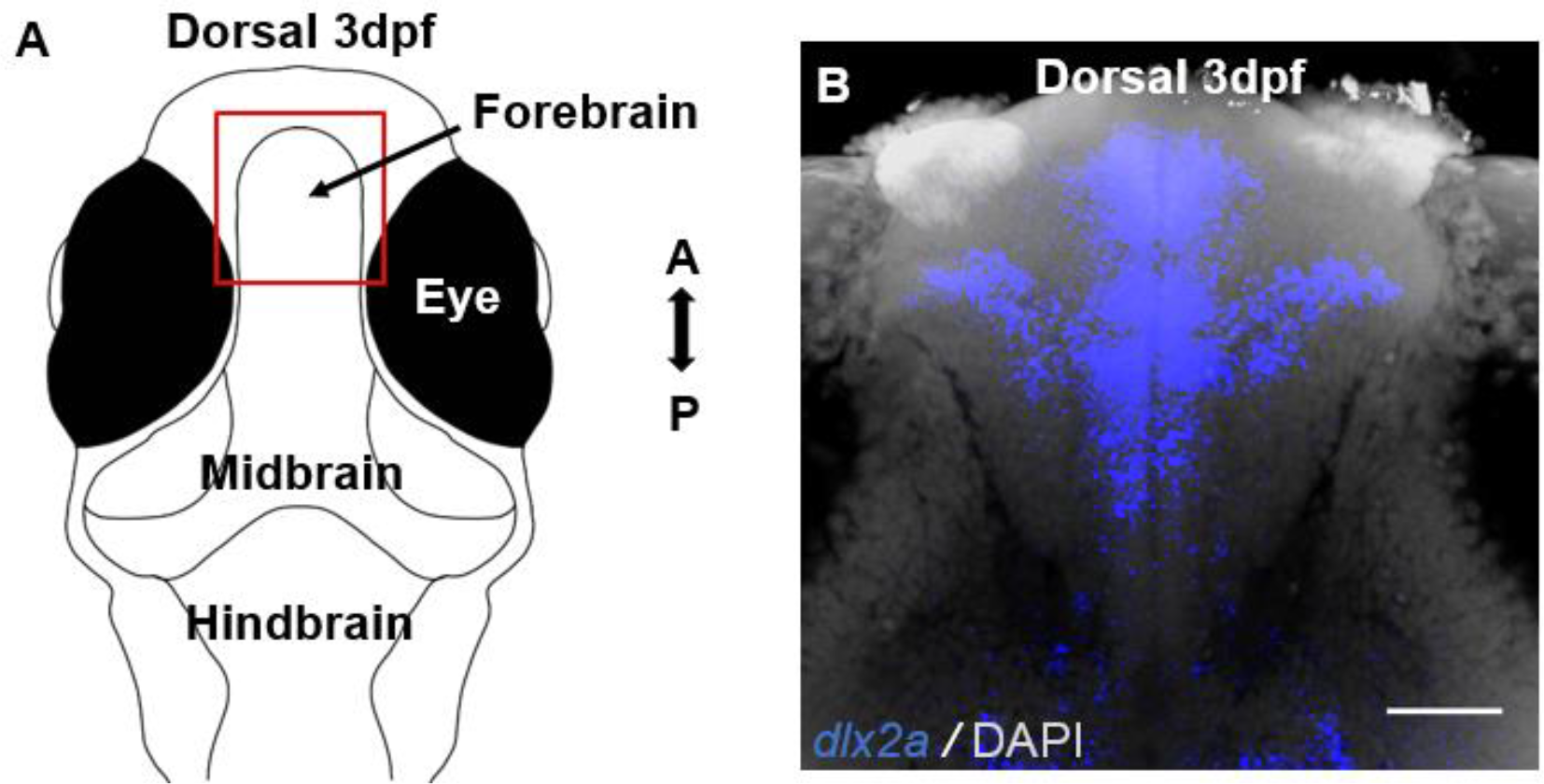
Dorsal view of *dlx2a* staining in the forebrain of 3 dpf larvae. **A)** Anatomical schematic outlining brain regions in 3 dpf fish. **B)** Z-projection of confocal sections rostral to the anterior commissure that contained *dlx2a* positive neurons. The projection consists of 80 planes with an interplane spacing of 2 μm. A, anterior; P, posterior. Scale bar = 50 μm.

**Figure 3.1.2:**
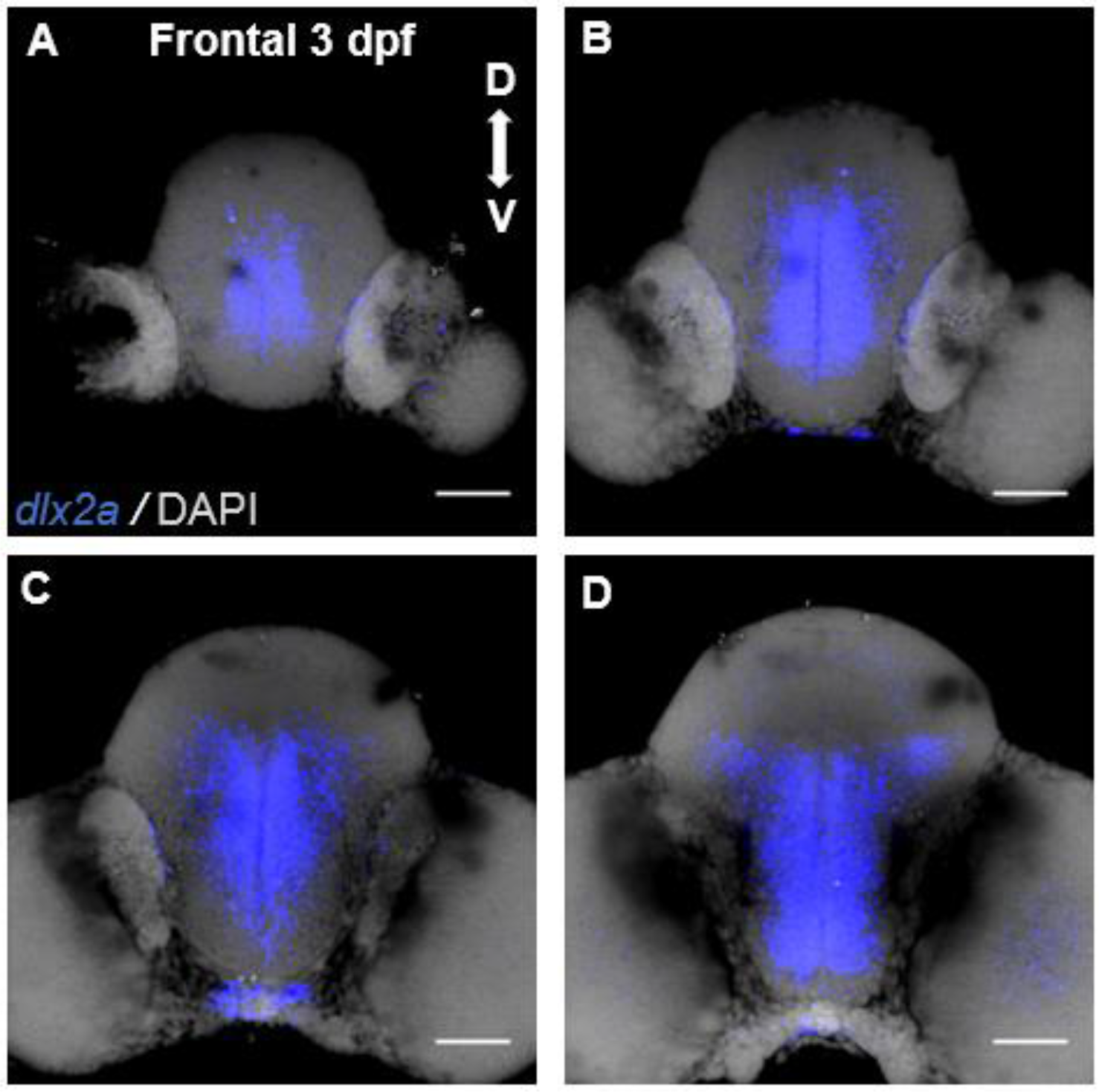
Frontal view of *dlx2a* in the forebrain of 3 dpf larvae. Each image is a confocal z-projection spanning 20μm. Images are organized from anterior to posterior, with **A** being the most anterior section. D, dorsal; V, ventral. Scale bar = 50 μm.

Mueller *et al.,* (2006) showed that the vast majority of neurons within the zebrafish subpallium are GABAergic. To identify GABAergic neurons within the subpallium across developmental stages (3, 7, 14, and 21 dpf), we performed whole mount immunostaining using the transgenic line *gad1b:|R|-GFP* (Satou *et al.,* 2013). At 3 dpf (n = 3), there was strong staining within the olfactory bulb and the subpallium with minimal GABA expression in the pallium (Figure 3.1.4, column A; Figure 3.1.3, A). At 7 dpf (n = 3) there was also strong olfactory bulb and subpallium staining (Figure 3.1.4, column B, panels 1-2; Figure 3.1.3, B). At this stage, GABA expression could be seen in the early migrated telencephalic region (Figure 3.1.4, column B, panels 5-7). 14 and 21 dpf larvae (n = 3) had a similar staining pattern within the olfactory bulb, subpallial, and pallial areas (Figure 3.1.4, column C; Figure 3.1.3, C). At 21 dpf (n = 3) subpallial GABAergic staining expanded to match the increase in brain volume of the subpallium. Staining within the pallium was observed in areas rostral to the anterior commissure (Figure 3.1.4, column D; Figure 3.1.3, D). This staining revealed a dense GABAergic population throughout the subpallium both rostral and caudal to the anterior commissure and is consistent with previously published expression data done in brain sections (Mueller, 2006).

**Figure 3.1.3:**
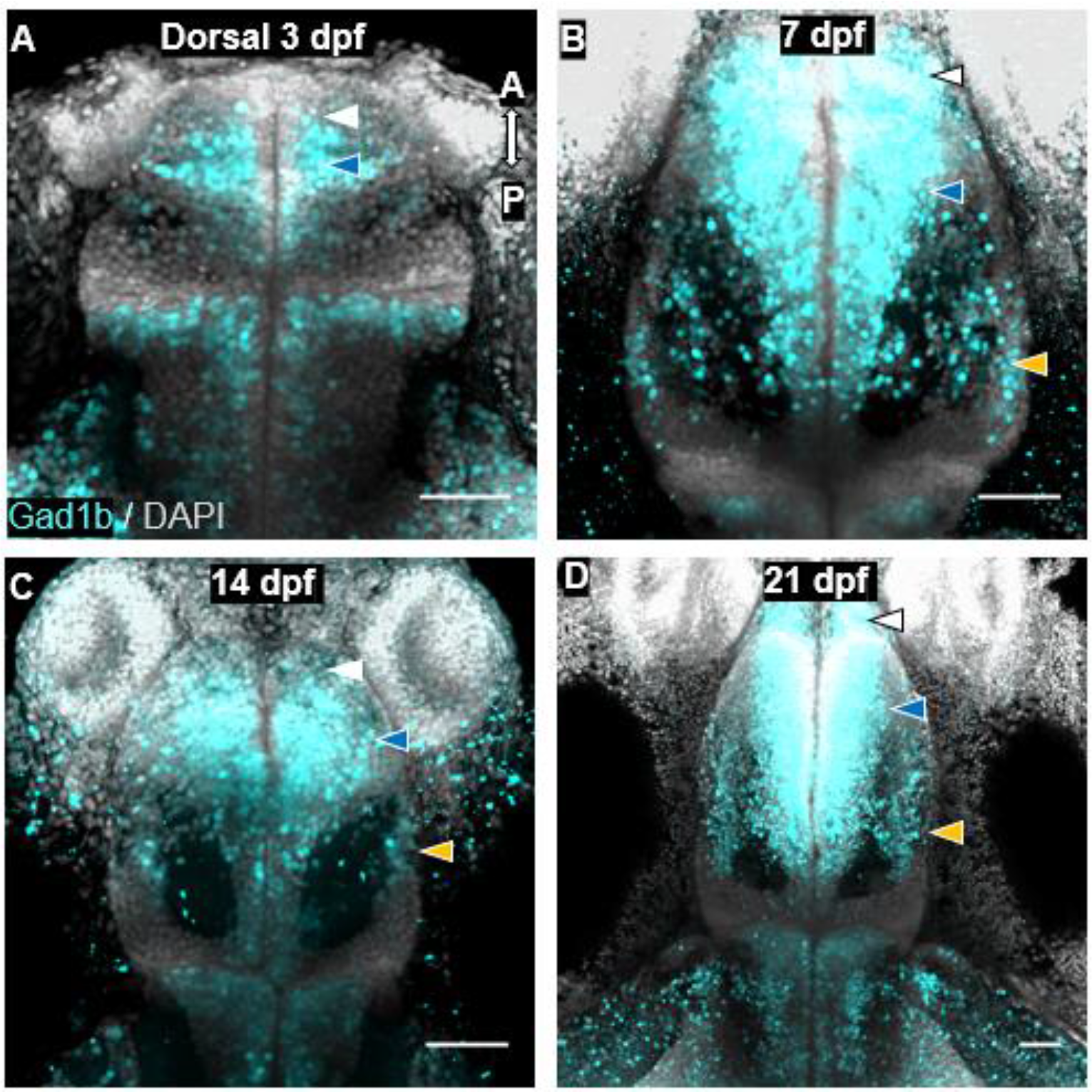
Dorsal view of GABAergic staining in the forebrain of developing larvae. Confocal z-projections of the forebrain of 3 dpf **(A)**, 7 dpf **(B),** 14 dpf **(C**), and 21 dpf **(D)** larvae. GABAergic staining is present in the olfactory bulb, precommissural subpallium, and lateral pallial areas. White arrowheads indicate staining in the olfactory bulb. Blue arrowheads indicate staining in the subpallium. Orange arrowheads indicate staining in the pallium. A, anterior; P, posterior. Scale bar = 50 μm.

**Figure 3.1.4:**
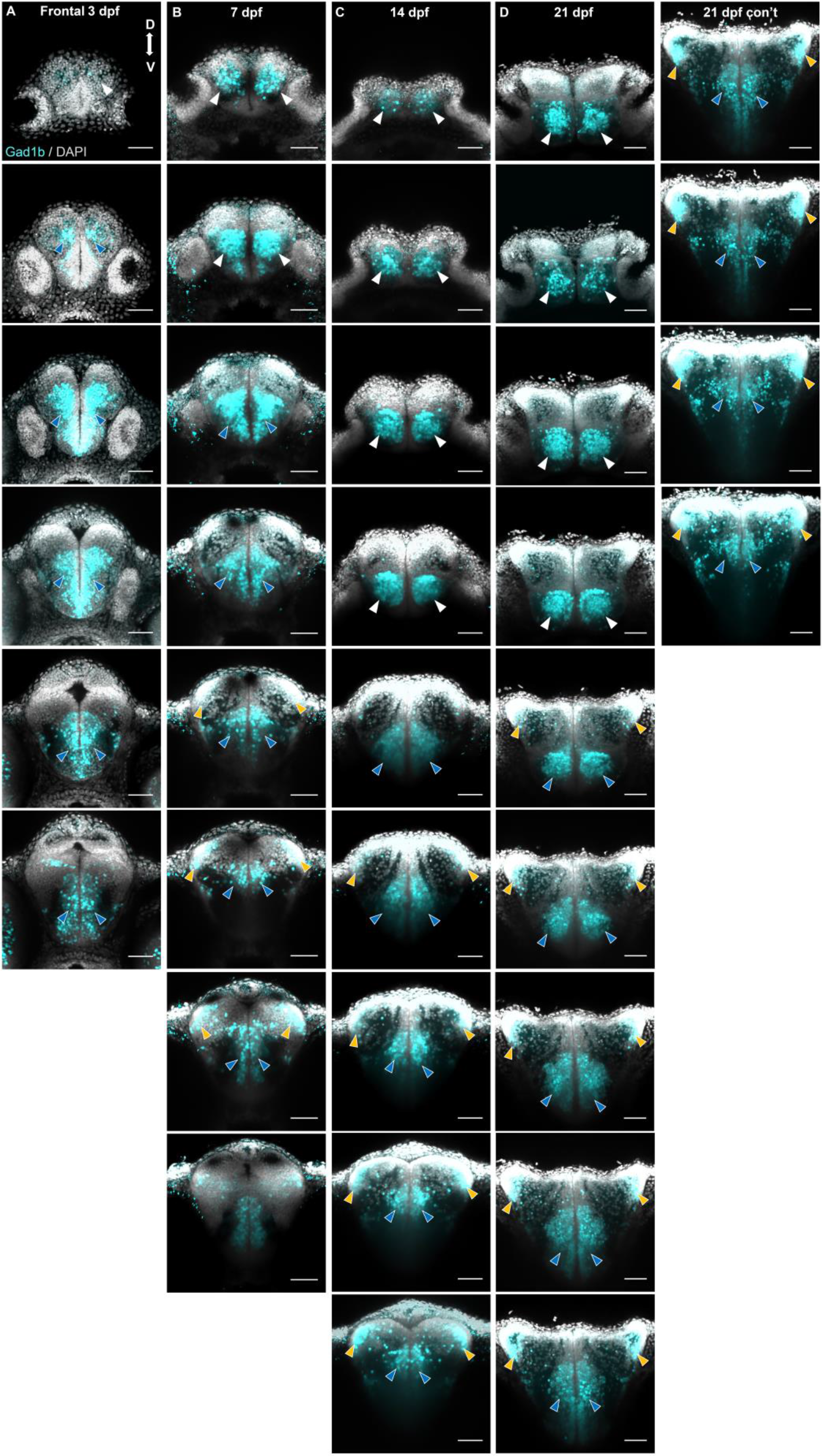
Frontal view of GABAergic neurons in the subpallium of 3, 7, 14, and 21 dpf larvae. Confocal z-projections of the forebrain of 3 dpf **(A)**, 7 dpf **(B),** 14 dpf **(C**), and 21 dpf **(D)** larvae. Sections are organized from anterior to posterior, with the most anterior sections in the top row (**A, B, C, D**). White arrowheads indicate staining in the olfactory bulb. Blue arrowheads indicate staining in the subpallium. Orange arrowheads indicate staining in the pallium. d, doral; v, ventral. Scale bar = 50 μm.

### 3.2 Development of Putative Direct Pathway Markers

#### 3.2.1 *tac1* Expression

Direct pathway neurons of the striatum express D1 receptors and use substance P as a neurotransmitter (Gerfen, 1992). To identify direct pathway neurons, we created RNA *in situ* probes for *tachykinin 1* (*tac1*), a precursor peptide that is cleaved to produce substance P. *In situ* processed samples differed slightly anatomically from the immunostained samples due to the heating required. We performed staining at 3, 7, 14, and 21 dpf. At 3 dpf, dense bilateral clusters of neurons (>100 cells) in close proximity to the olfactory bulbs were present, and no staining was observed near the anterior commissure (Figure 3.2.1, A; Figure 3.2.2, column C, panels 1-3). An average of 6.4 neurons (n=5, two-tailed t-test, p < 0.0001) were found dorsal and rostral to the anterior commissure at 7 dpf. There was also robust expression near or within the olfactory bulbs (>200 neurons) as well as bilateral staining close to the midline pallial/ subpallial boundary (Figure 3.2.1, B; Figure 3.2.2, column B, panels 2-4). At 14 dpf, an average of 10.3 *tac1* positive neurons (n=5, two-tailed t-test, p < 0.0001) were present just dorsal and rostral to the anterior commissure (Figure 3.2.1, C; Figure 3.2.2, column C, panels 4-5). The dense staining at the olfactory bulb impeded reliable cell estimates at stages beyond 7 dpf. There was bilateral staining in the medial dorsal telencephalic area, but it is unclear whether cell bodies were stained (Figure 3.2.2, column D, panels 7-8). The precommissural *tac1* neuronal population increased to an average of 20.4 neurons (n=5, two-tailed t-test, p < 0.001) at 21 dpf (Figure 3.2.1, D; Figure 3.2.2, column D, panels 6-8). The majority of *tac1* neurons were confined dorsally and rostrally to the anterior commissure, with 1-2 neurons appearing in the ventral subpallium. The robust staining at the level of the olfactory bulb impeded reliable cell counts at this stage. There was bilateral staining of the medial dorsal telencephalic area present, however clear cell bodies were not visible. At each stage, the number of *tac1* neurons increased (Table 3.4.1, Graph 3.2.1).

**Figure 3.2.1:**
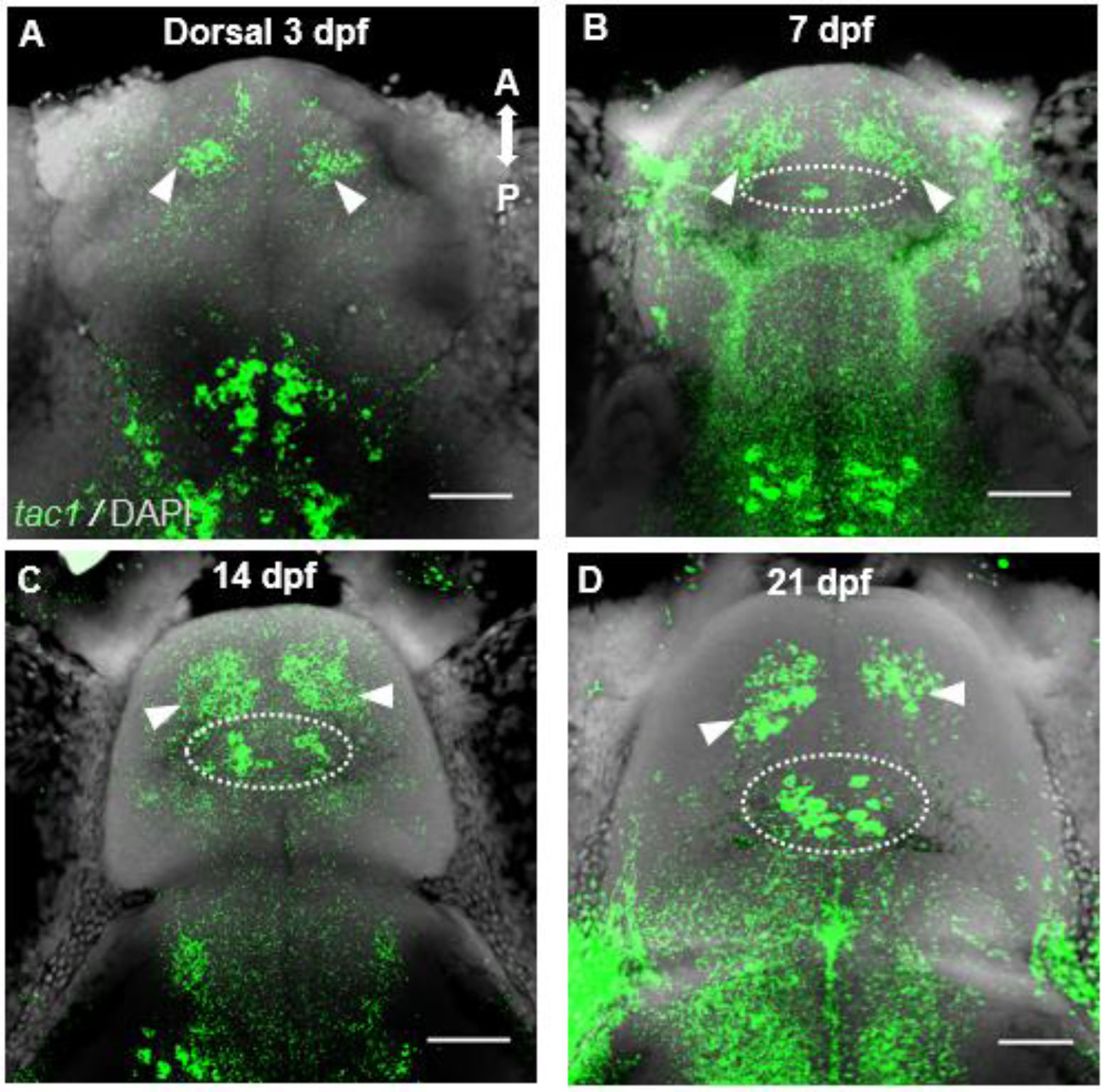
Dorsal view of *tac1* expression in the forebrain during larval development. Confocal z-projections of the forebrain from 3 dpf **(A)**, 7 dpf **(B),** 14 dpf **(C**), and 21 dpf **(D)** larvae. *tac1* staining is present in the olfactory bulbs (white arrowheads) and precommissural subpallium (white hashed circle). A, anterior; P, posterior. Scale bar = 50 μm.

**Figure 3.2.2:**
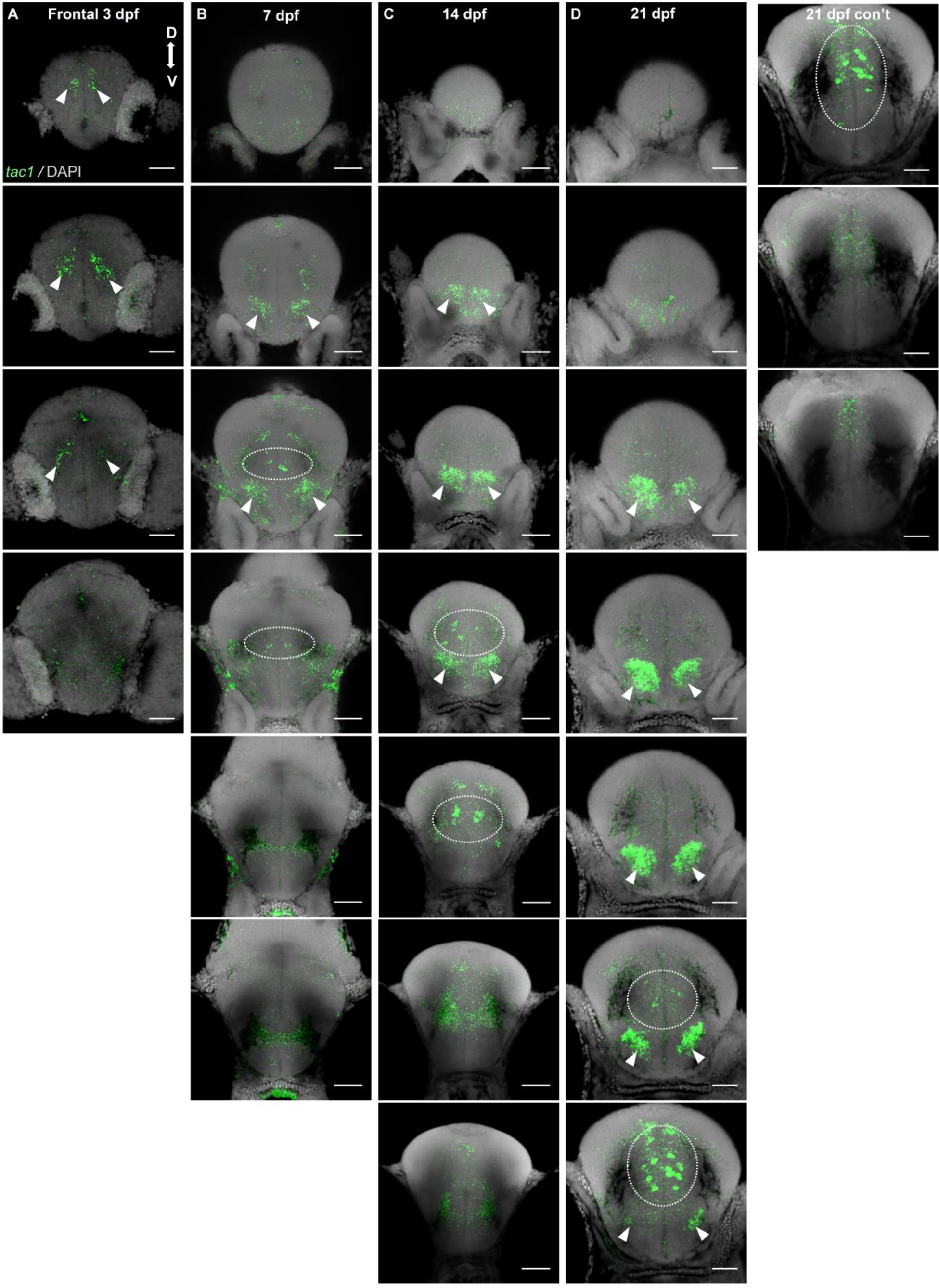
Frontal view of *tac1* expression in the subpallium of 3, 7, 14, and 21 dpf larvae. Confocal z-projections (20 µm) from the forebrain from 3 dpf **(A)**, 7 dpf **(B),** 14 dpf **(C**), and 21 dpf **(D)** larvae. Sections are organized from anterior to posterior, with the most anterior sections in the top row **(A, B, C, D)**. *tac1* staining is present in the olfactory bulbs (white arrowheads) and precommissural subpallium (white hashed circle). D, dorsal; V, ventral Scale bar = 50 μm.

**Graph 3.2.1:**
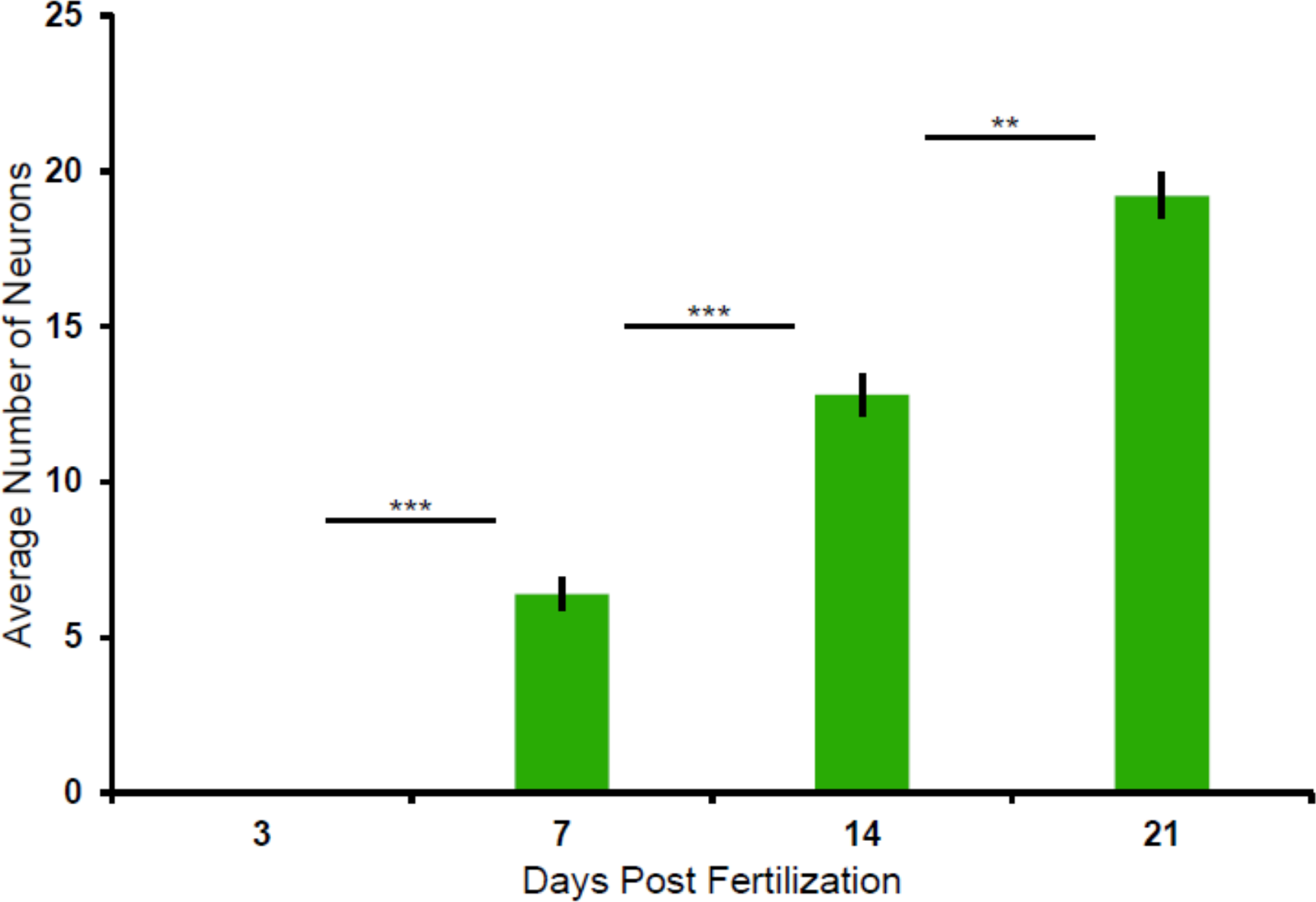
Cell counts for the precommissural subpallial *tac1* population. Two-tailed unpaired t-tests were performed, with statistically significant results were reported as p < 0.01 with the Bonferroni correction. *** p < 0.0001. ** p < 0.001 Data are represented as mean ± SEM.

A linear regression model was applied and the resulting R^2^ value was 0.98, indicating a strong positive linear correlation between age and the number of subpallial *tac1* neurons (Graph 3.2.2). To validate the penetration of our *tac1 in situ* probe in our whole mount samples, we also performed FISH in dissected brains (n = 3) for 7 and 21 dpf larvae. There was no difference in the number of labelled neurons in the two preparations indicating our whole mount samples accurately represent *tac1* expression (Table 3.3.1, Graph 3.3.3, two-tailed t-tests, p > 0.05).

**Graph 3.2.2:**
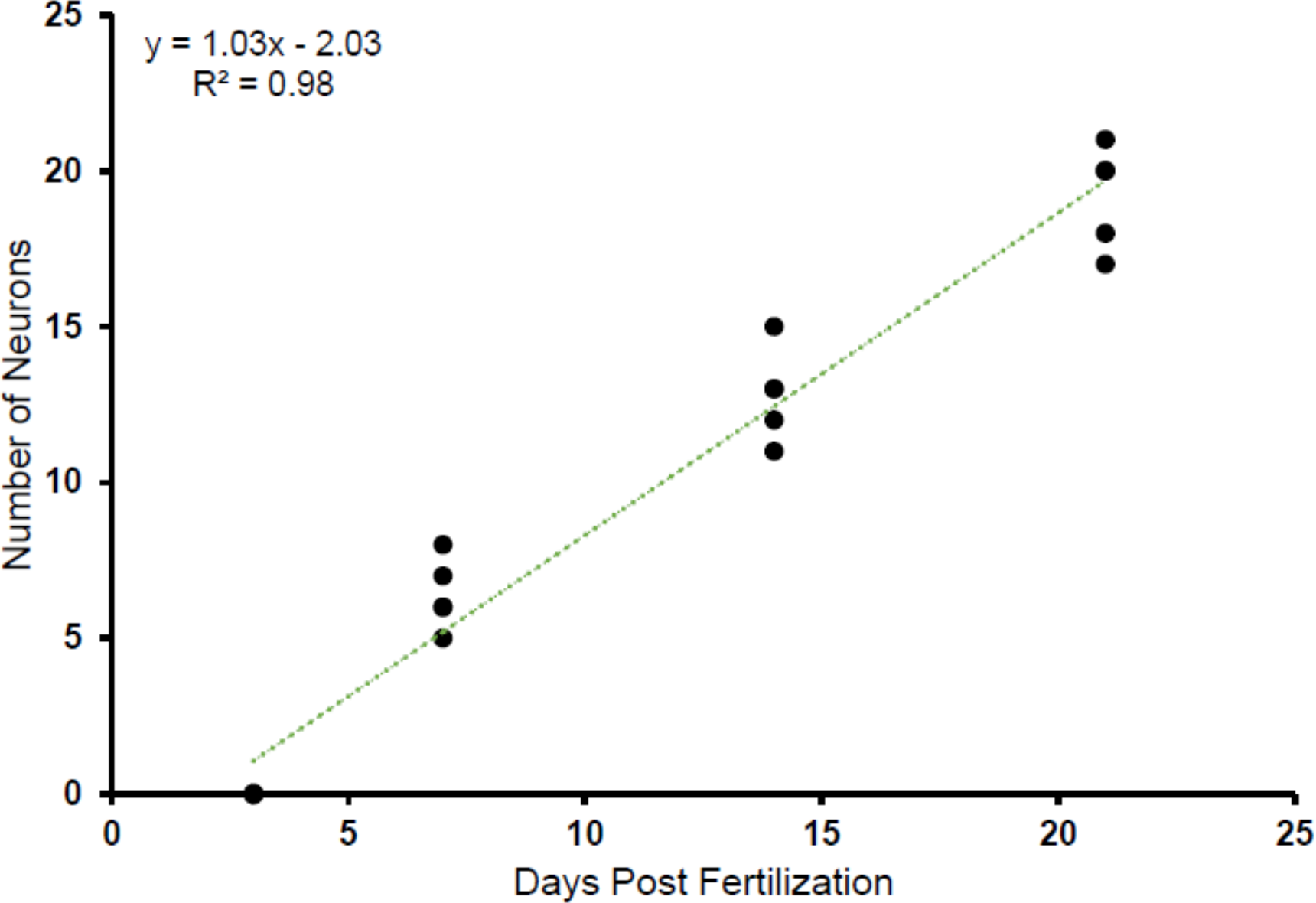
The precommissural subpallial *tac1* neuronal population increases linearly with age. Scatterplot showing *tac1* neuron counts in the precommissural subpallium at 3, 7, 14, and 21 dpf larvae. R^2^ = 0.98.

**Graph 3.3.3:**
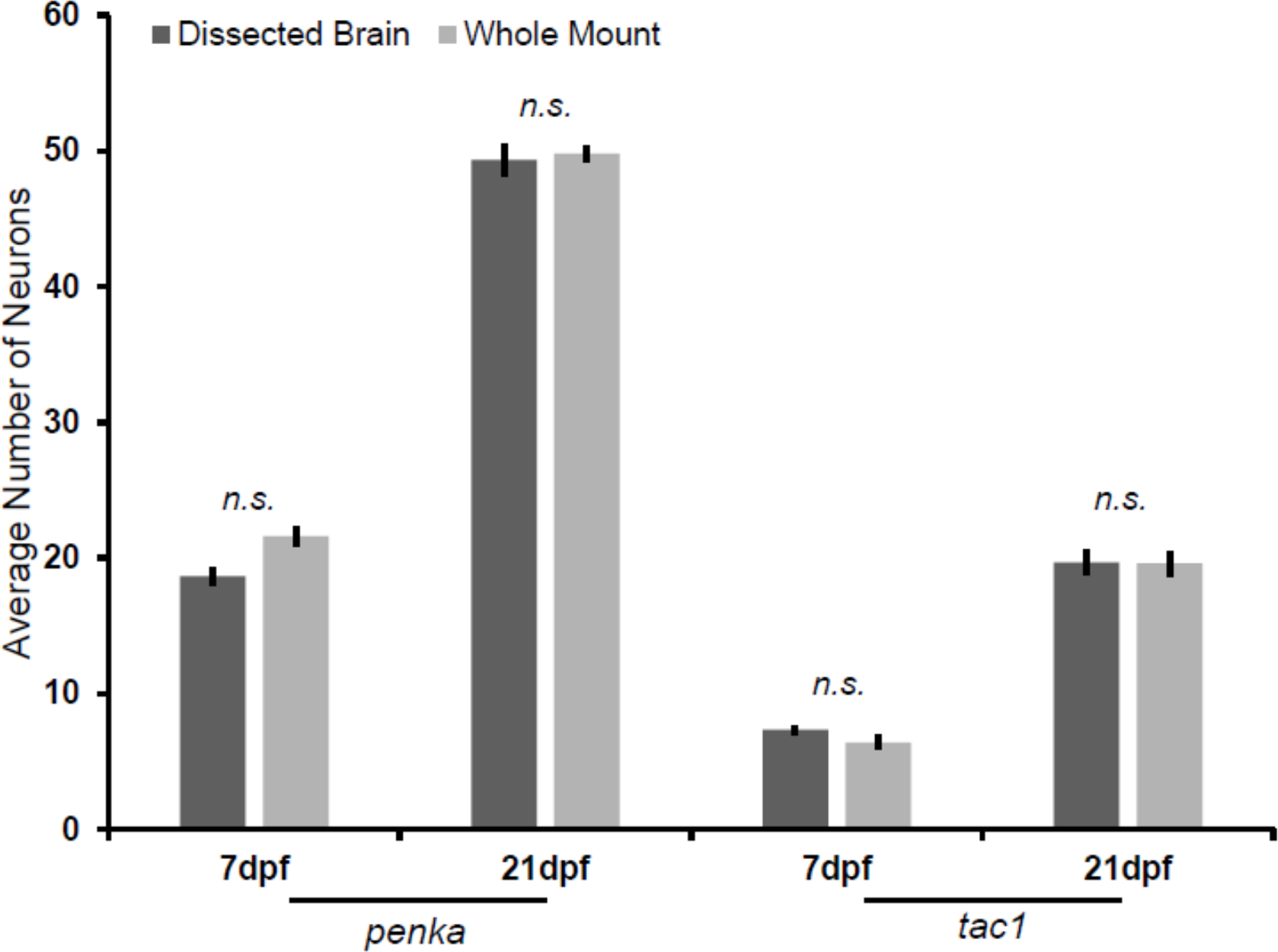
Comparison between *tac1* and *penka* cell counts in dissected brains and whole mount samples. Fluorescent *in situ* hybridization in whole mount (n = 5) dissected brains (n = 3) results in the same number of labelled neurons for *penka* and *tac1*. Two-tailed unpaired t-tests were performed, and Bonferroni corrected. n.s., not significant. Data are represented as mean ± SEM.

**Table 3.3.1:**
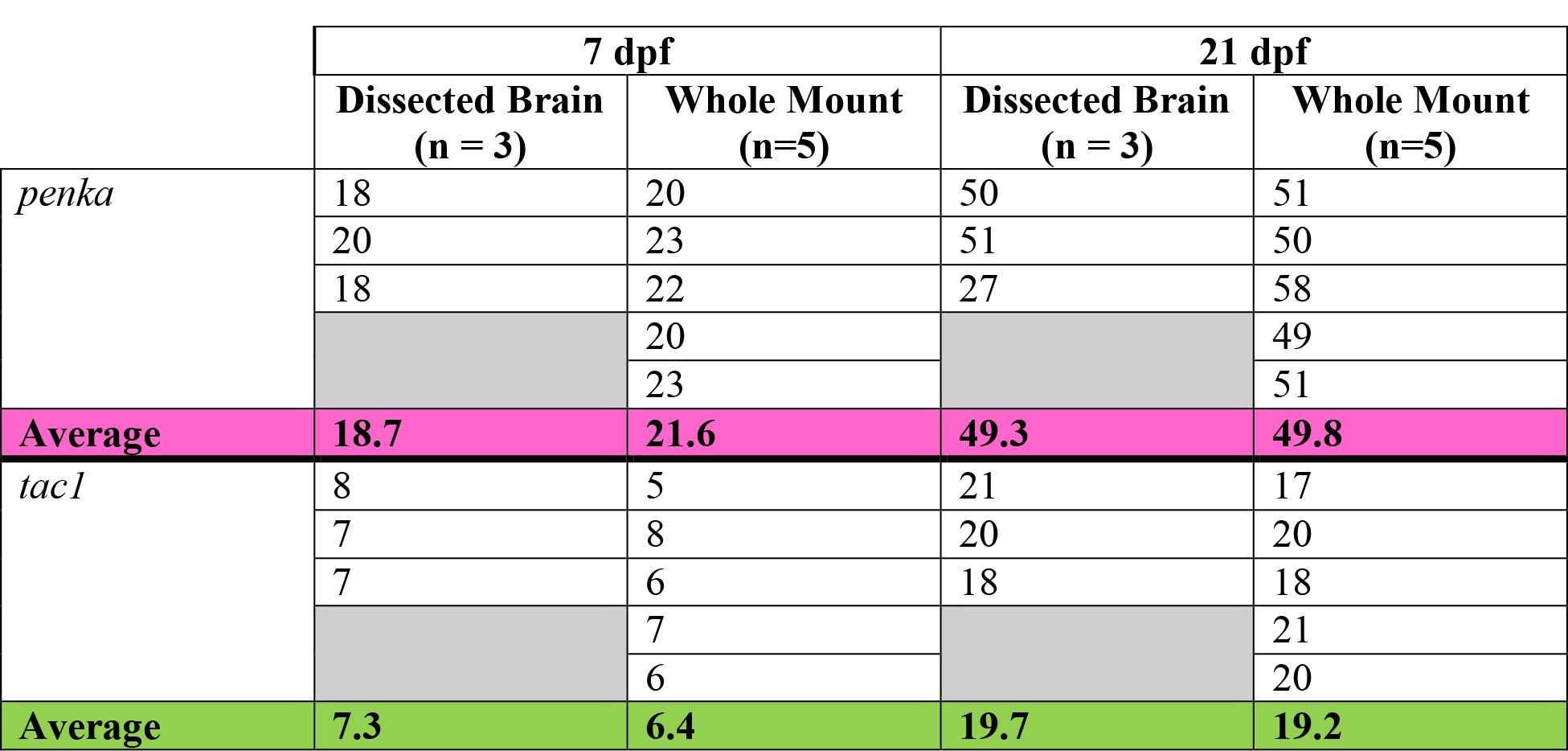
Average number of *tac1* and *penka* positive neurons in dissected brains and whole mount samples.

|  | 7 dpf |  | 21 dpf |  |
| --- | --- | --- | --- | --- |
|  | Dissected Brain<br>(n = 3) | Whole Mount<br>(n=5) | Dissected Brain<br>(n = 3) | Whole Mount<br>(n=5) |
| *penka* | 18 | 20 | 50 | 51 |
|  | 20 | 23 | 51 | 50 |
|  | 18 | 22 | 27 | 58 |
|  |  | 20 |  | 49 |
|  |  | 23 |  | 51 |
| **Average** | **18.7** | **21.6** | **49.3** | **49.8** |
| *tac1* | 8 | 5 | 21 | 17 |
|  | 7 | 8 | 20 | 20 |
|  | 7 | 6 | 18 | 18 |
|  |  | 7 |  | 21 |
|  |  | 6 |  | 20 |
| **Average** | **7.3** | **6.4** | **19.7** | **19.2** |

**Table 3.4.1:** Average number of *tac1* and *penka* positive neurons at different developmental timepoints.

|  | 3 dpf | 7 dpf | 14 dpf | 21 dpf |
| --- | --- | --- | --- | --- |
| *penka* | 2 | 20 | 39 | 51 |
|  | 5 | 23 | 37 | 50 |
|  | 3 | 22 | 37 | 58 |
|  | 2 | 20 | 31 | 49 |
|  | 3 | 23 | 32 | 51 |
| **Average** | **3** | **21.6** | **35.2** | **49.8** |
| *tac1* | 0 | 5 | 11 | 17 |
|  | 0 | 8 | 13 | 20 |
|  | 0 | 6 | 12 | 18 |
|  | 0 | 7 | 15 | 21 |
|  | 0 | 6 | 13 | 20 |
| **Average** | **0** | **6.4** | **12.8** | **19.2** |

#### 3.2.2 Substance P Expression

Since alternative splicing of *tac1* can lead to multiple neuropeptides, we used a substance P antibody to validate our *in situ* staining for 3, 7, 14, and 21 dpf larvae. Cell bodies were not clearly labelled with this antibody, so we are unable to correlate *in situ tac1* cell counts with substance P immunostaining. At 3 dpf, there was robust expression in the fibres of the developing olfactory bulb (Figure 3.2.3, A; Figure 3.2.4, column A, panels 1-2) that overlapped with staining observed in our *tac1 in situs*. The olfactory bulb fibres remained robustly stained in 7 dpf larvae (Figure 3.2.4, column B, panels 1-4). Substance P was also observed at this stage within fibres traversing through the anterior commissure, lateral forebrain bundles, habenula, and tectum (Figure 3.2.3, B). At 14 dpf, there was a dense Substance P positive network of fibres innervating the pallium, along with expression in the olfactory bulb, anterior commissure, lateral forebrain bundles, habenula, and tectum (Figure 3.2.3, C; Figure 3.2.4, column C). This expression pattern was similar in 21 dpf larvae (Figure 3.2.3, D; Figure 3.2.4, column D). At all timepoints, there was robust staining close to the border of the subpallium and the olfactory bulb in a similar location to the most rostral *tac1* neuronal population seen in our *in situs.* Staining extended from this border region to the densely innervate areas throughout the bulb. This staining pattern is consistent with the large, most rostral population of *tac1* neurons being part of the olfactory bulb (Figure 3.2.5).

**Figure 3.2.3:**
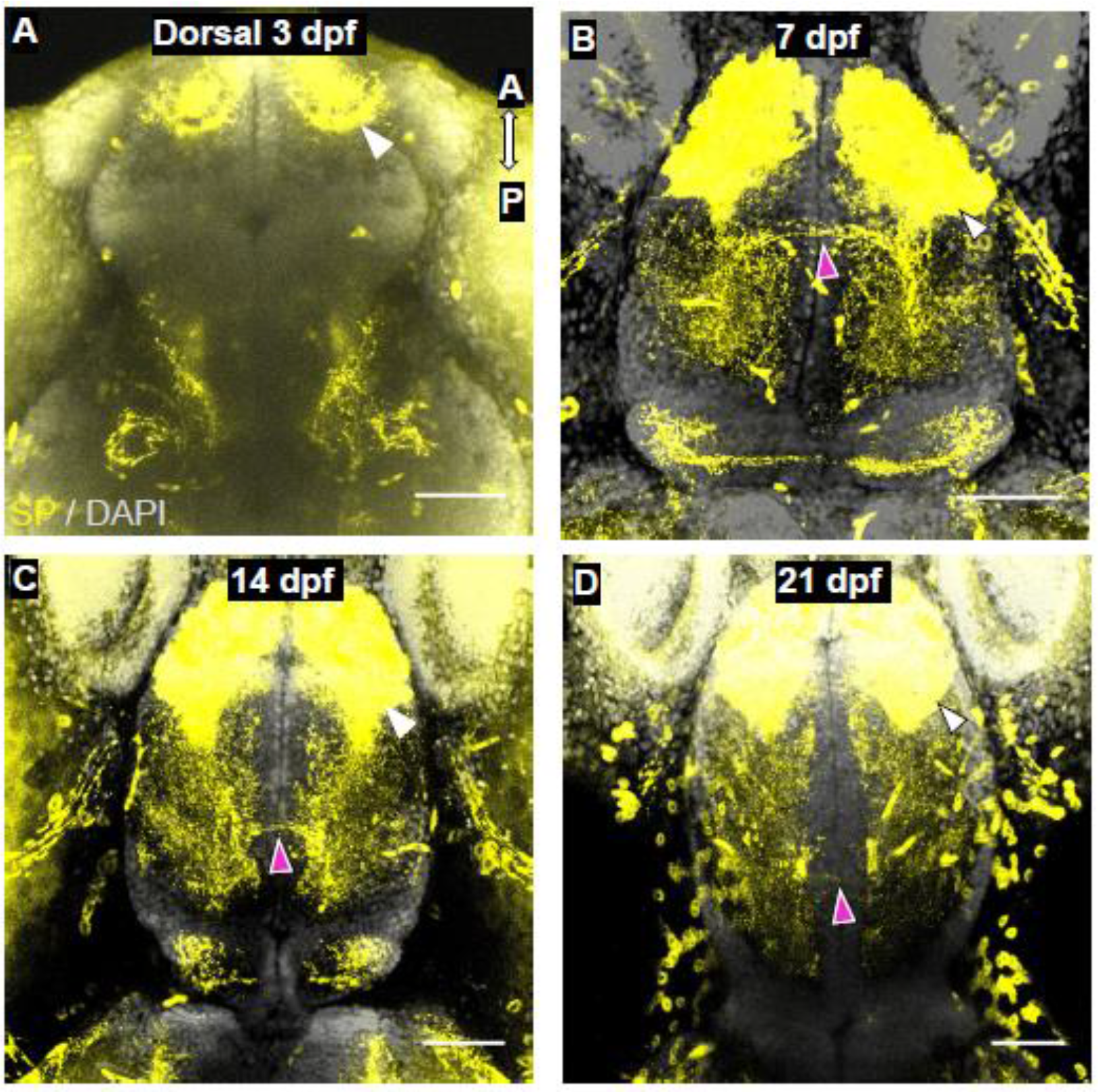
Dorsal view of substance P staining in the forebrain of developing larvae. Confocal z-projections of the forebrain of 3 dpf **(A)**, 7 dpf **(B),** 14 dpf **(C**), and 21 dpf **(D)** larvae. Substance P staining is present in the olfactory bulb, anterior commissure, and lateral forebrain bundles. White arrowheads indicate staining in the olfactory bulb. Purple arrowheads indicate fibre staining within tracts and the anterior commissure. A, anterior; P, posterior. Scale bar = 50 μm.

**Figure 3.2.4:**
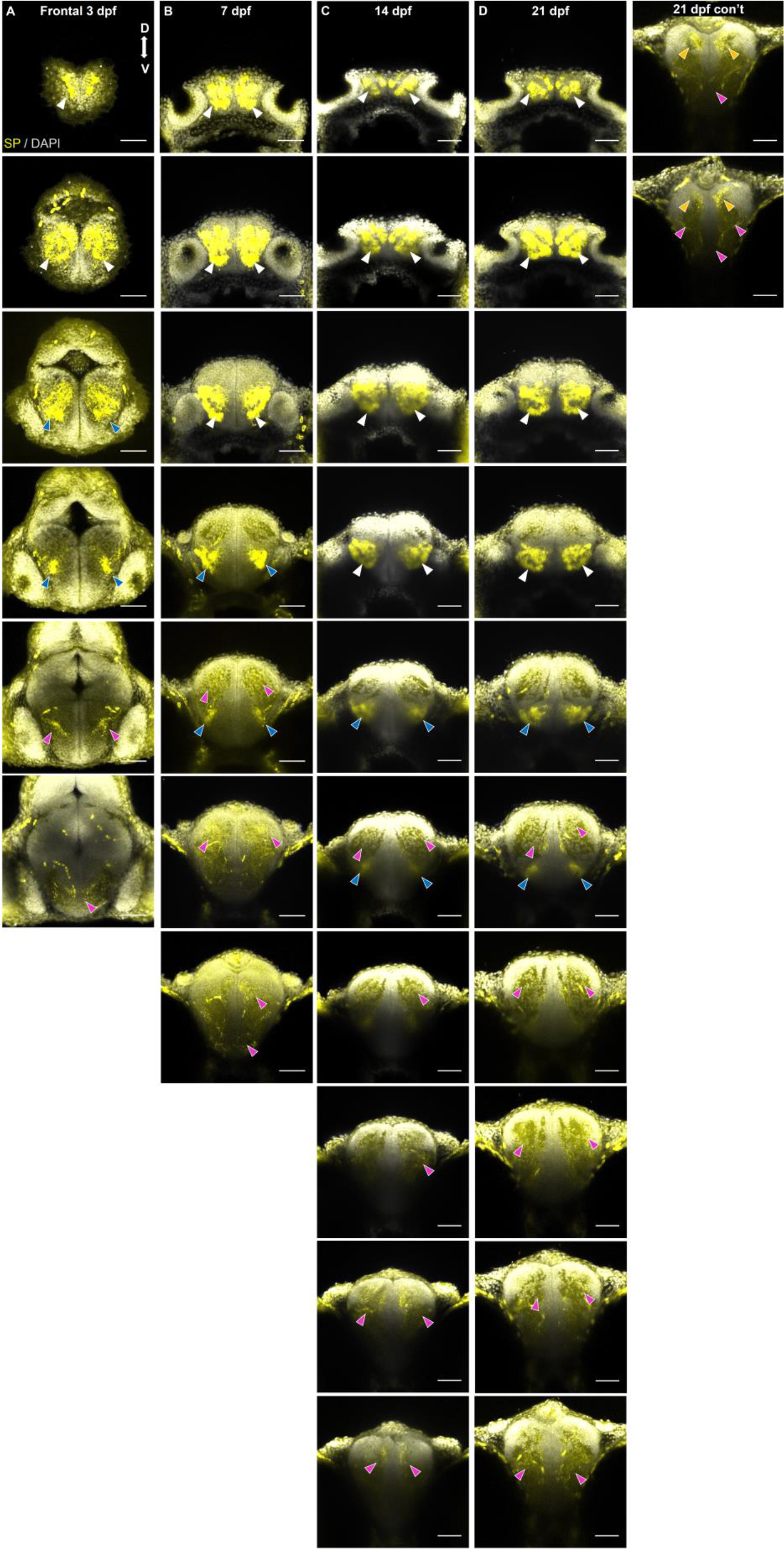
Frontal view of substance P neurons in the subpallium of 3, 7, 14, and 21 dpf larvae. Confocal z-projections (20 µm) of the forebrain of 3 dpf **(A)**, 7 dpf **(B),** 14 dpf **(C**), and 21 dpf **(D)** larvae. Sections are organized from anterior to posterior, with the most anterior sections in the top row (**A, B, C, D**). White arrowheads indicate staining in the olfactory bulb. Purple arrowheads indicate staining of the fibres within the tracts and the anterior commissure. Blue arrowheads indicate staining in the subpallium. Orange arrowheads indicate pallial staining. Surface background fluorescence was due to the need to increase laser power in order to detect substance P staining. D, dorsal; V, ventral. Scale bar = 50 μm.

**Figure 3.2.5:**
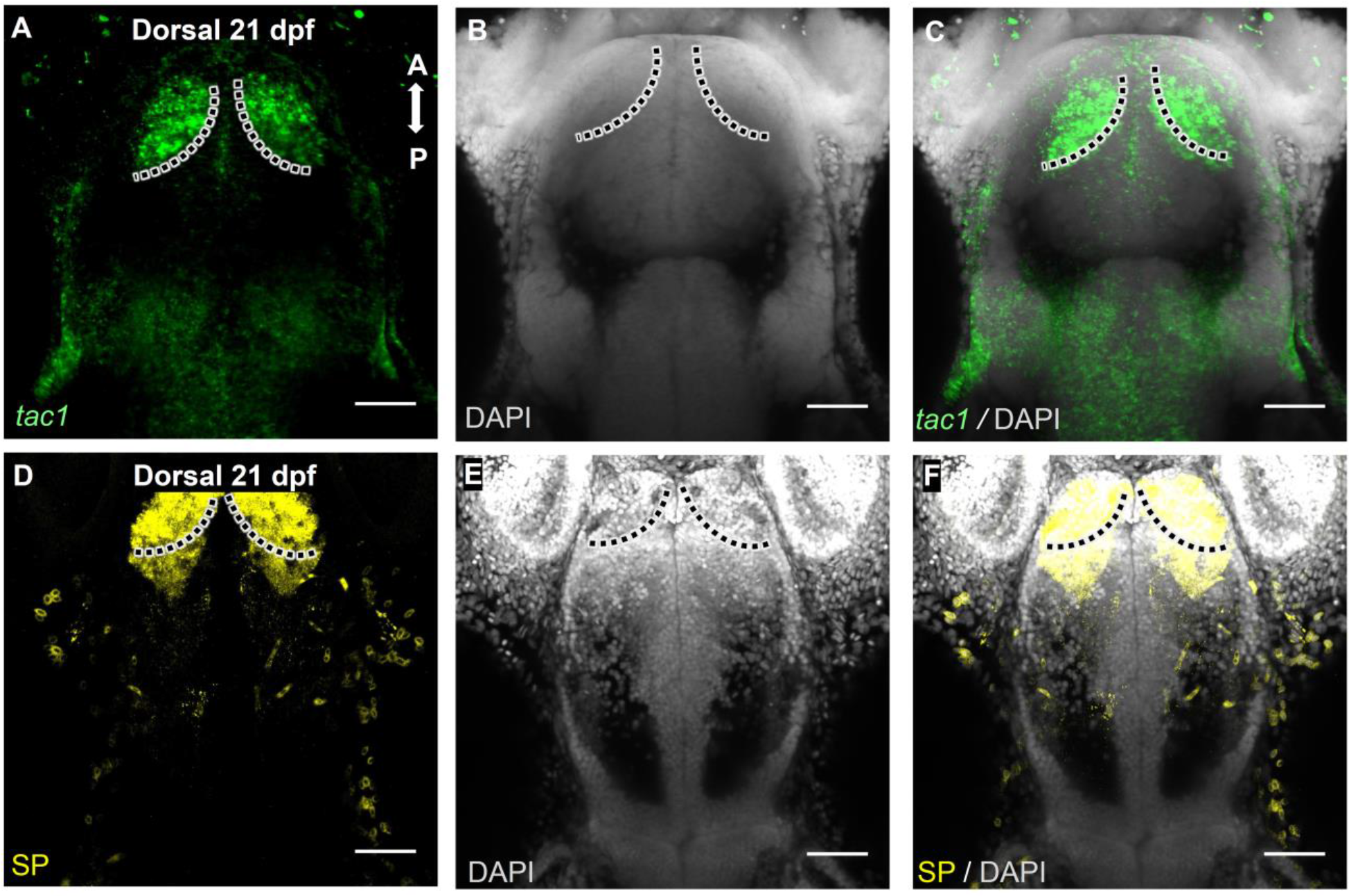
Dorsal view of *tac1* and substance P expression within the olfactory bulb. Sections of 21 dpf larvae are shown as z-projections representing planes that have olfactory bulb staining present. Each row is colour coded for a different marker: *tac1* (**A-C)** and substance P **(D-F)**. A, anterior; P, posterior. Scale bar = 50 μm.

### 3.3 Development of Putative Indirect Pathway Markers

#### 3.3.1 *penka* Expression

Indirect pathway neurons within the striatum express D2 receptors and use enkephalin as a neurotransmitter (Gerfen, 1992). We created RNA *in situ* probes for proenkephalin A (*penka)*, the precursor peptide to enkephalin, in order to visualize the expression pattern of potential indirect pathway neurons at 3, 7, 14, and 21 dpf. At 3 dpf, there was an average of 3 neurons (n = 5) in the precommissural subpallial area, with the majority of *penka* neurons at this stage situated in preoptic areas caudal to the anterior commissure (Figure 3.3.1, A; Figure 3.3.2, column A). This population increased to an average of 22.6 neurons (n = 5, two-tailed t-test, p < 0.0001) at 7 dpf. These neurons extended in a column-like fashion throughout the dorsoventral axis of the subpallium just rostral to the anterior commissure (Figure 3.3.1, B; Figure 3.3.2, column B, panels 3-5). At 14 dpf, the precommissural population increased to an average of 35.2 neurons (n = 5, two-tailed t-test, p < 0.0001) which extended along the midline along the dorsoventral subpallial axis (Figure 3.3.1, C; Figure 3.3.2, column C, panels 6-8). The *penka* population increased at 21 dpf, averaging 49.8 neurons (n = 5, two-tailed t-test, p < 0.0001) with robust staining just dorsal and near the level of the anterior commissure, extending ventrally (Figure 3.3.1, D; Figure 3.3.2, column D, panels 7-11). At each stage, the number of *penka* neurons increased (Table 3.4.1, Graph 3.3.1). A linear regression model was applied and the resulting R^2^ value was 0.96, indicating a strong linear correlation between age and the number of subpallial *penka* neurons (Graph 3.3.2). We also performed FISH on dissected brains (n = 3) to validate the penetration of our *penka in situ* probe in our whole mount samples for 7 and 21 dpf larvae. There was no significant difference between the number of *penka* neurons in both preparations, indicating that our whole mount samples accurately represent *penka* expression (Table 3.3.1, Graph 3.3.3, two-tailed t-tests, p > 0.05).

**Figure 3.3.1:**
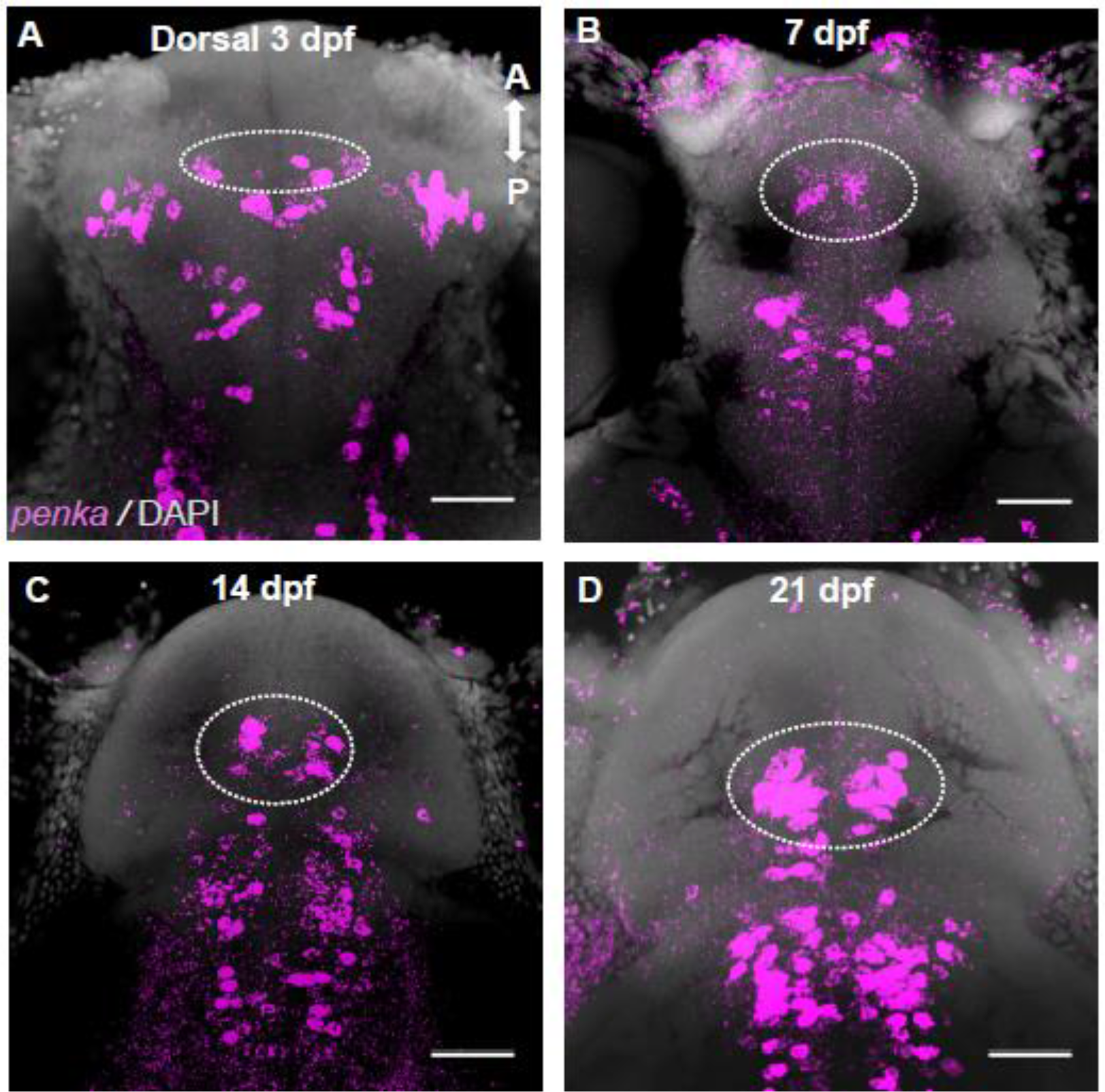
Dorsal view of *penka* expression in the forebrain during larval development. Sections 3 dpf **(A)**, 7 dpf **(B),** 14 dpf **(C**), and 21 dpf **(D)** larvae are shown as z-projections representing planes that have neurons present. *penka* staining is present in the precomissural area (denoted by white hashed circle), telencephalon, and rostral midbrain structures. A, anterior; P, posterior. Scale bar = 50 μm.

**Figure 3.3.2:**
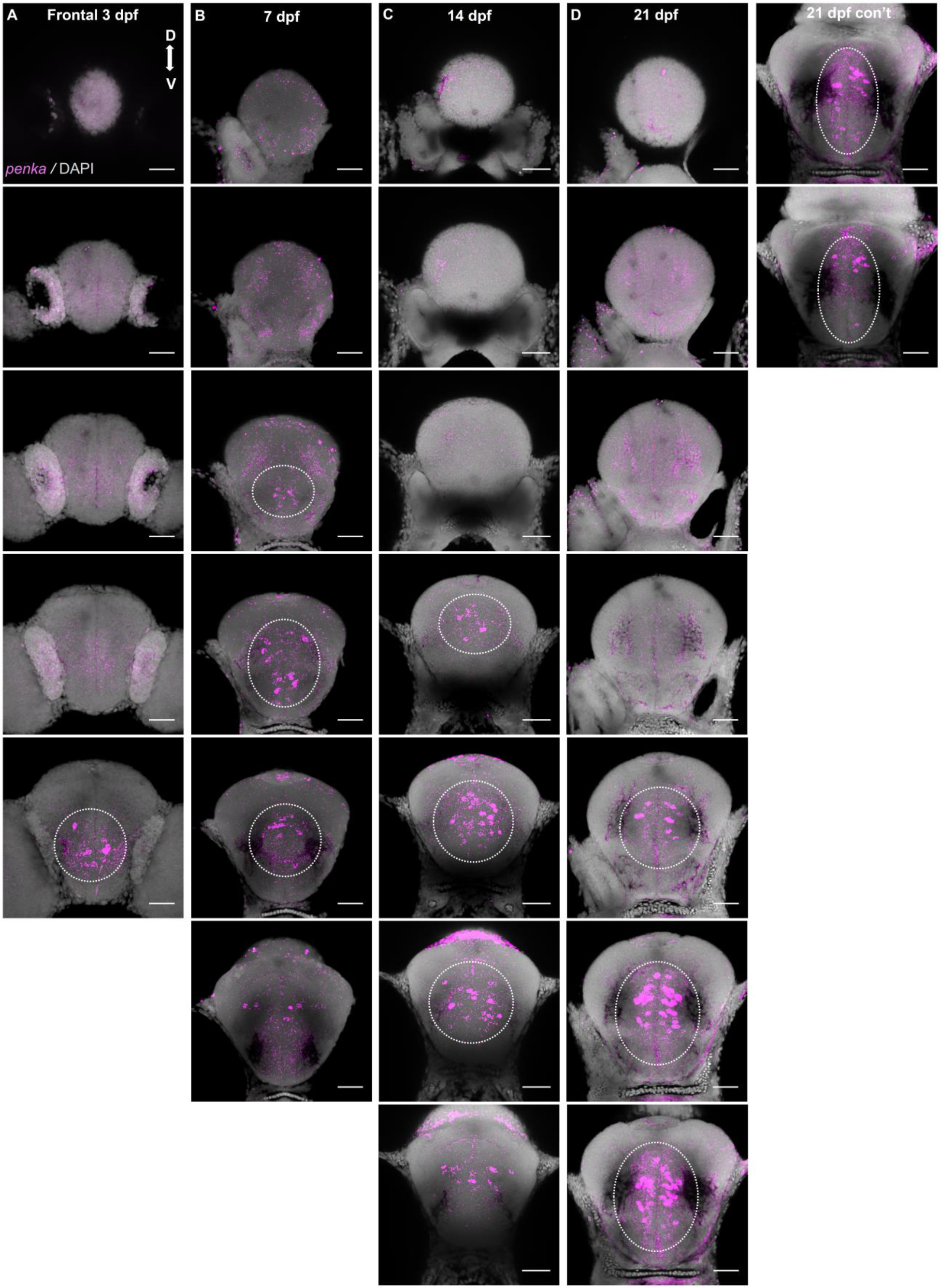
Frontal view of *penka* expression in the subpallium of 3, 7, 14, and 21 dpf larvae. Confocal z-projections (20 µm) from the forebrain from 3 dpf **(A)**, 7 dpf **(B),** 14 dpf **(C**), and 21 dpf **(D)** larvae. Sections are organized from anterior to posterior, with the most anterior sections in the top row **(A, B, C, D)**. *penka* staining is present in the precommissural subpallium (white hashed circle). D, dorsal; V, ventral. Scale bar = 50 μm.

**Graph 3.3.1:**
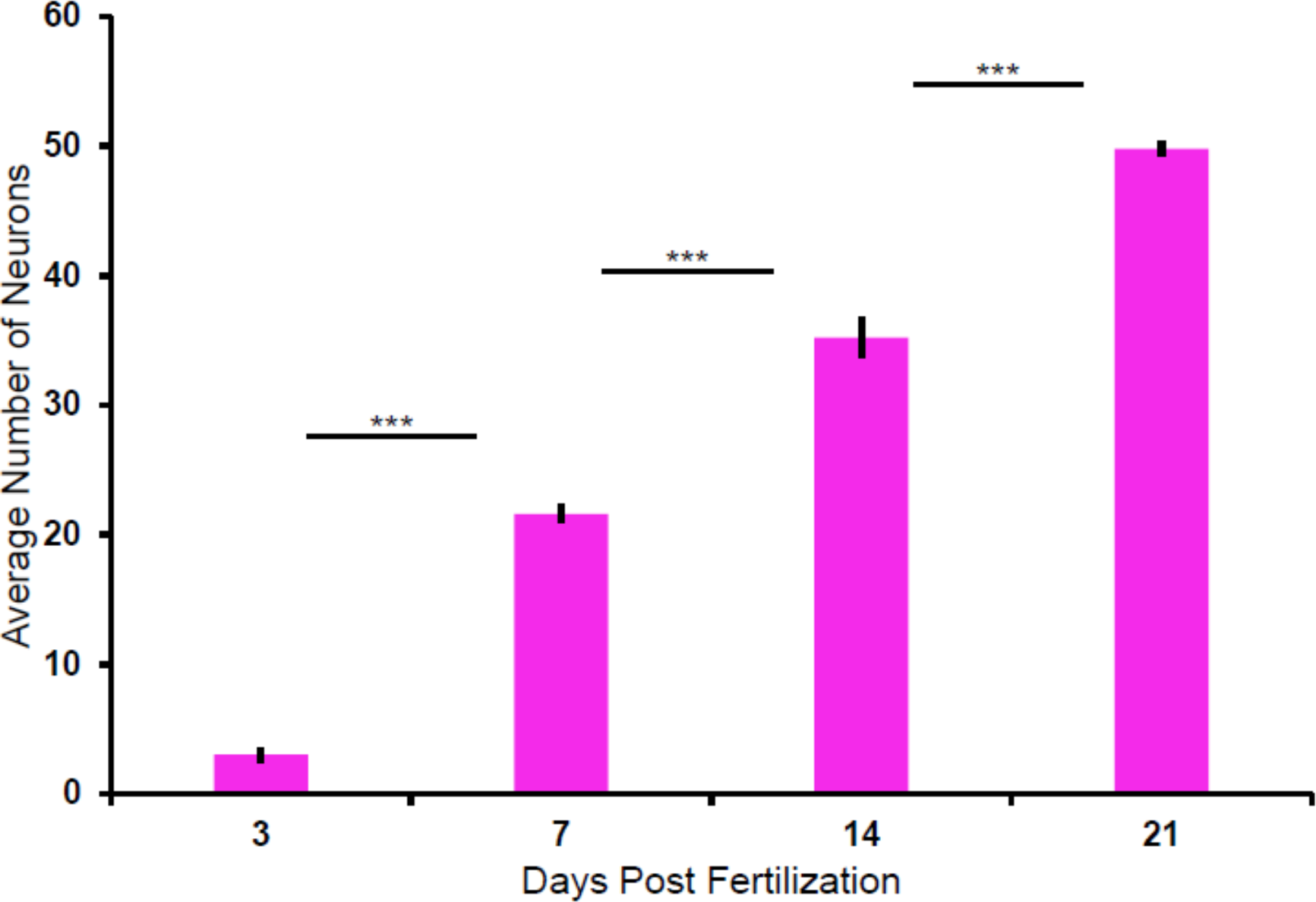
Cell counts for the precommissural subpallial *penka* population. Two-tailed unpaired t-tests were performed, with statistically significant results were reported as p < 0.01 with Bonferroni correction. *** p < 0.0001. Data are represented as mean ± SEM.

**Graph 3.3.2:**
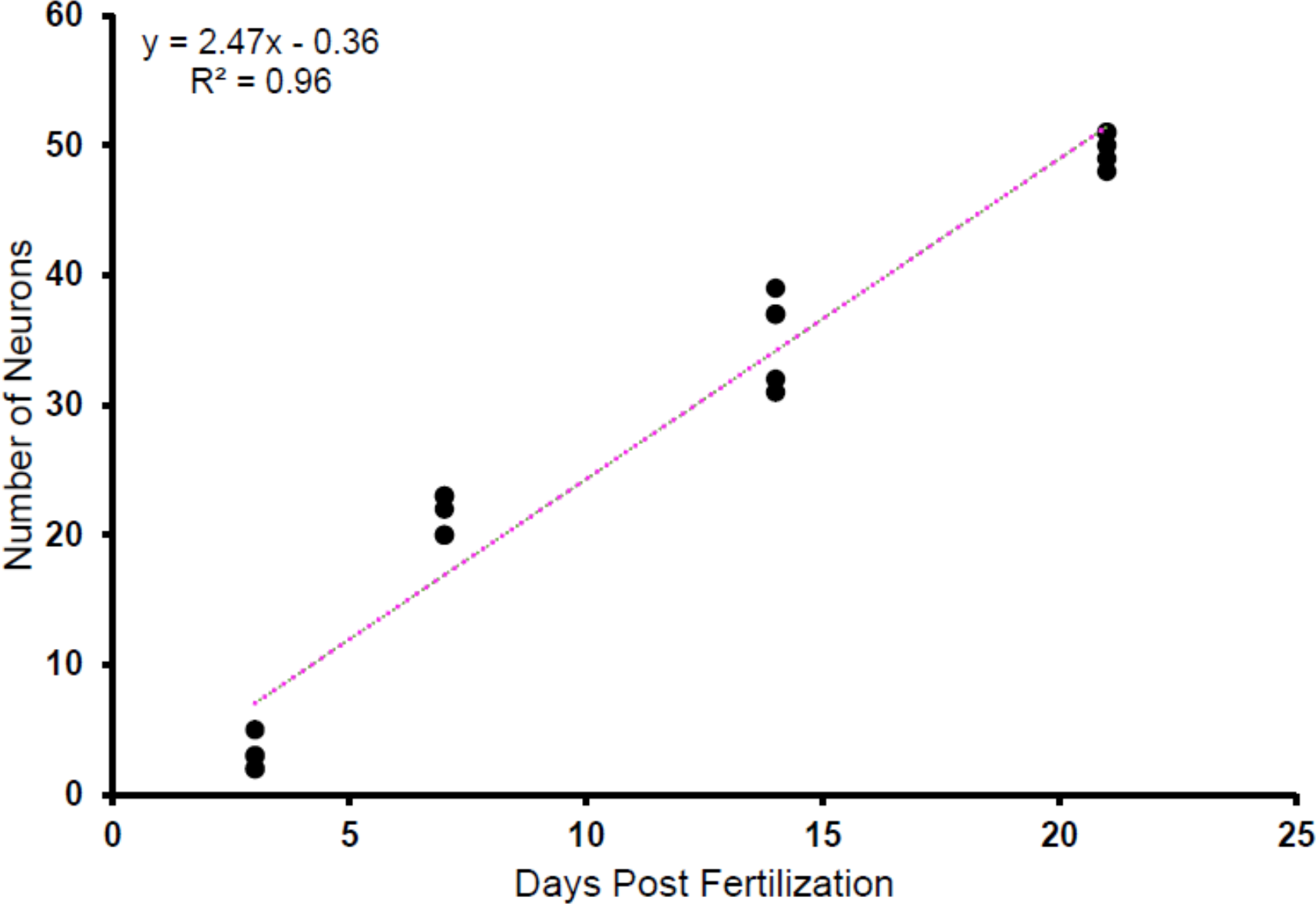
The precommissural subpallial *penka* neuronal population increases linearly with age. Scatterplot showing the total count of *penka* neurons in the precommissural subpallium at 3, 7, 14, and 21 dpf larvae. R^2^ = 0.96.

#### 3.3.2 Enkephalin Expression

To further validate the expression pattern of *penka*, we used an enkephalin antibody at 3, 7, 14, and 21 dpf. At 3 dpf (n = 3), there was dense staining of fibres within the olfactory bulb, anterior commissure, and lateral forebrain bundles (Figure 3.3.4, column A). There was also bilateral staining within the early migrated telencephalic regions (Figure 3.3.3, A). Staining within the olfactory bulb, anterior commissure, and lateral forebrain bundles persisted at 7 dpf (n = 3), with fibres extending into pallial zones (Figure 3.3.3, B; Figure 3.3.4, column B). Potential cell bodies were labelled at the midline within the dorsal subpallium rostral to the anterior commissure (Figure 3.3.4, column B, panels 3-5). At 14 dpf (n = 3), there were potential cell bodies labelled dorsal and rostral to the anterior commissure which extended ventrally along the midline (Figure 3.3.4, column C, panels 4-5). Robust staining of the olfactory bulb fibres, anterior commissure, lateral forebrain bundles, and tracts within the pallium were also present (Figure 3.3.3, C).

**Figure 3.3.3:**
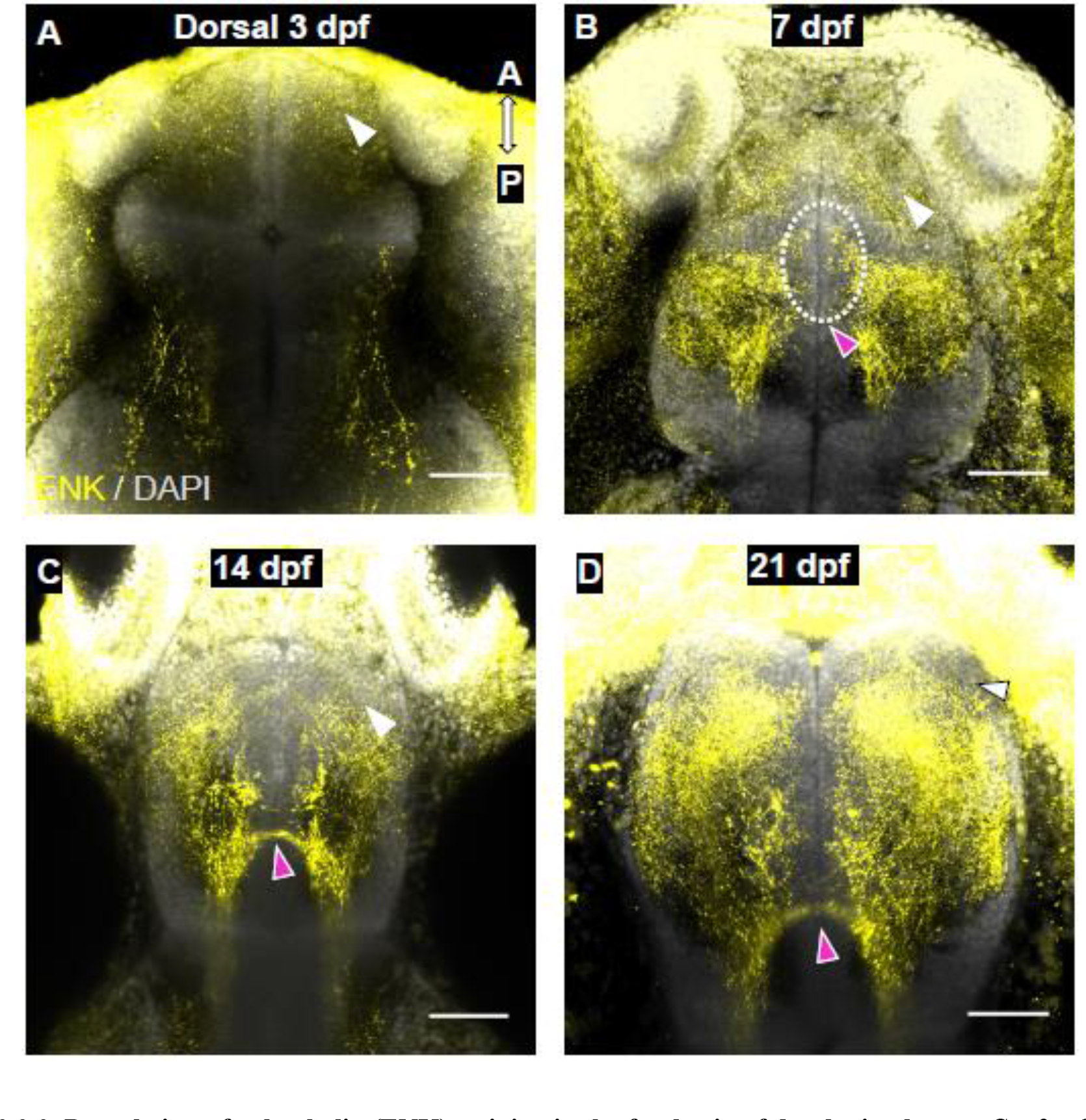
Dorsal view of enkephalin (ENK) staining in the forebrain of developing larvae. Confocal z-projections of the forebrain of 3 dpf **(A)**, 7 dpf **(B),** 14 dpf **(C**), and 21 dpf **(D)** larvae. GABAergic staining is present in the olfactory bulb, precommissural subpallium, and lateral forebrain bundles. White arrowheads indicate staining in the olfactory bulb. Purple arrowheads indicate staining of the fibres within the tracts and the anterior commissure. White hashed circle indicates clusters of enkephalin neurons. Surface background fluorescence was due to the need to increase laser power in order to detect enkephalin staining. A, anterior; P, posterior. Scale bar = 50 μm.

**Figure 3.3.4:**
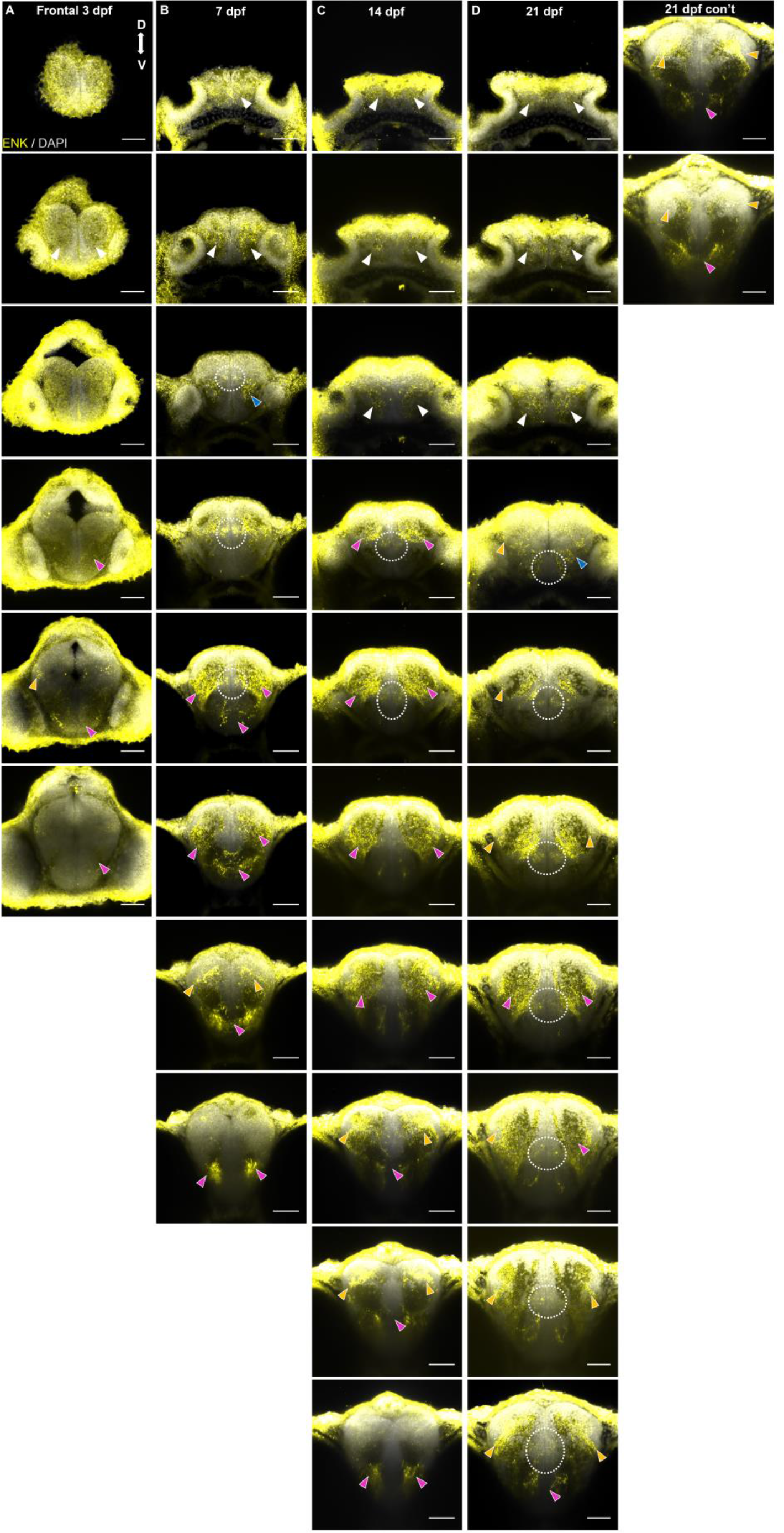
Frontal view of enkephalin neurons in the subpallium of 3, 7, 14, and 21 dpf larvae. Confocal z-projections (20 µm) of the forebrain of 3 dpf **(A)**, 7 dpf **(B),** 14 dpf **(C**), and 21 dpf **(D)** larvae. Sections are organized from anterior to posterior, with the most anterior sections in the top row (**A, B, C, D**). White arrowheads indicate staining in the olfactory bulb. Purple arrowheads indicate staining of the fibres within the tracts and the anterior commissure. Blue arrowheads indicate staining in the subpallium. Orange arrowheads indicate pallial staining. White hashed circle indicates clusters of enkephalin neurons. Surface background fluorescence was due to the need to increase laser power in order to detect enkephalin staining. D, dorsal; V, ventral. Scale bar = 50μm.

Similar staining was found in 21 dpf larvae (n = 3) alongside an increase in potentially labelled cell bodies grouped within the midline of the dorsal subpallium (Figure 3.3.3, D; Figure 3.3.4, column D, panels 4-10). The robust staining of fibres within each timepoint made cell counting unreliable and we were unable to correlate the number of enkephalin labelled neurons to our *penka in situ* labelled neuronal population.

### 3.4 Comparison of *tac1* and *penka* Neuronal Populations at Different Larval Stages

At each timepoint, there were significantly more *penka* neurons than *tac1* neurons within the precommissural subpallium (Table 3.4.1, Graph 3.4.1). A majority of the *tac1* neuronal population remained within a distinct volume dorsal and rostral to the anterior commissure (Figure 3.4.1, A-D, I-L). *penka* neurons were found to expand along the dorsoventral axis at 7 dpf onward (Figure 3.4.1, E-H, M-P). To determine *tac1* and *penka* expression relative to each other, we performed double fluorescent *in situ* hybridization at 7, 14, and 21 dpf. In 7 dpf, *tac1* and *penka* neurons were comingled dorsal to the anterior commissure, with the majority of the *penka* population found in more ventral regions of the subpallium (Figure 3.4.2, A; Figure 3.4.3 column A). The volume of space where both *tac1* and *penka* neurons were expressed was 48 x 39 x 31μm (x, y, z) in dimension and stained neurons started to appear ∼46μm away from the anterior most point of the larvae (Table 3.4.3, Graph 3.4.2). At 14 dpf, *tac1* and *penka* populations were also comingled in the dorsal precommissural subpallial region, with *penka* neurons extending along the midline into the ventral areas of the subpallium (Figure 3.4.2, B; Figure 3.4.3, column B). The volume of space where both markers were present was 68 x 46 x 44μm (x, y, z) in dimension and stained neurons started to appear ∼60μm from the anterior most point (Table 3.4.3, Graph 3.4.3). At 21 dpf, there was extensive overlap between *tac1* and *penka* neurons in the dorsal subpallium at dorsal precommissural levels with the majority of *penka* neurons extending ventrally along the midline (Figure 3.4.2, C; Figure 3.4.3, column C). Both markers were confined to a 68 x 49 x 54μm volume within the subpallium and stained neurons started to appear ∼84μm caudal to the anterior most point of the forebrain (Table 3.4.3, Graph 3.4.4). We counted the number of *penka* neurons within the *tac1* volume as an estimate for the number of *penka* neurons in the putative striatal region. At 7 dpf, the average number of *tac1* and *penkA* neurons within the *tac1* boundary were 5 and 4, respectively (n = 2). This increased to 12 and 10 in 14 dpf larvae (n = 2), and in 21 dpf larvae we counted an average of 19.5 *tac1* neurons (n = 2) and 20 *penka* neurons (n = 2) (Table 3.4.2).

**Figure 3.4.1:**
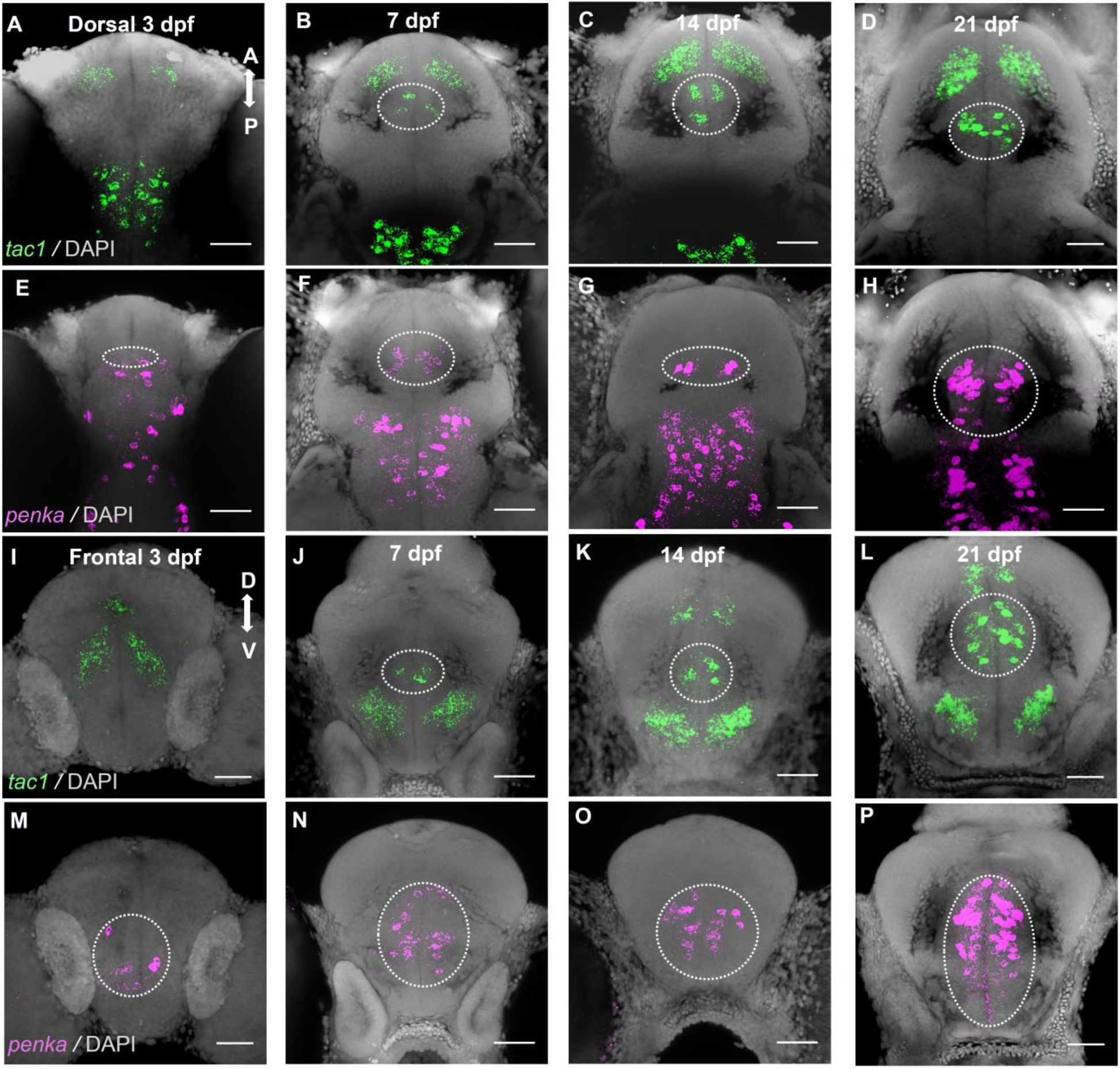
Comparison of *tac1* and *penka* expression across larval development. Dorsal views of the forebrain of 3 dpf **(A, E)**, 7 dpf **(B, F)**, 14 dpf **(C, G)**, 21 dpf **(D, H)** dpf larvae shown as confocal z-projections. Frontal views are confocal z-projections (20 µm) from the forebrain of 3 dpf **(I, M)**, 7 dpf **(J, N)**, 14 dpf **(K, O)**, 21 dpf **(L, P)** dpf larvae. White dashed circle indicates precommissural subpallial *tac1* and *penka* neurons at comparable sections. A, anterior; P, posterior; D, dorsal; V, ventral. Scale bar = 50μm.

**Figure 3.4.2:**
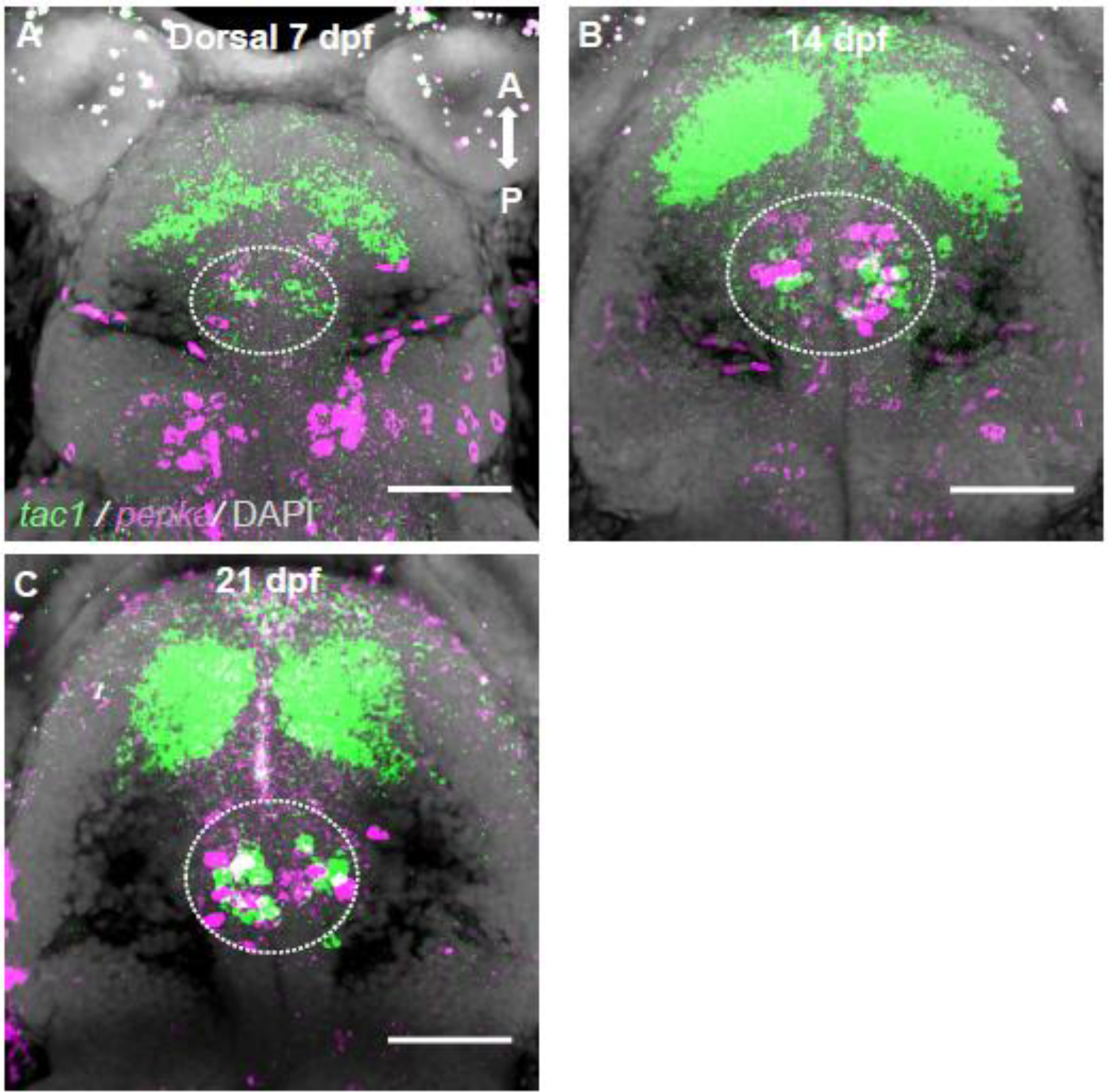
Dorsal view of double fluorescent *in situs* for *tac1* and *penka* during larval development. Confocal z-projections of the forebrain from 3 dpf **(A)**, 7 dpf **(B),** 14 dpf **(C**), and 21 dpf **(D)** larvae. *tac1* and *penka* are both present in the precommissural subpallium (white hashed circle). A, anterior; P, posterior. Scale bar = 50μm.

**Figure 3.4.3:**
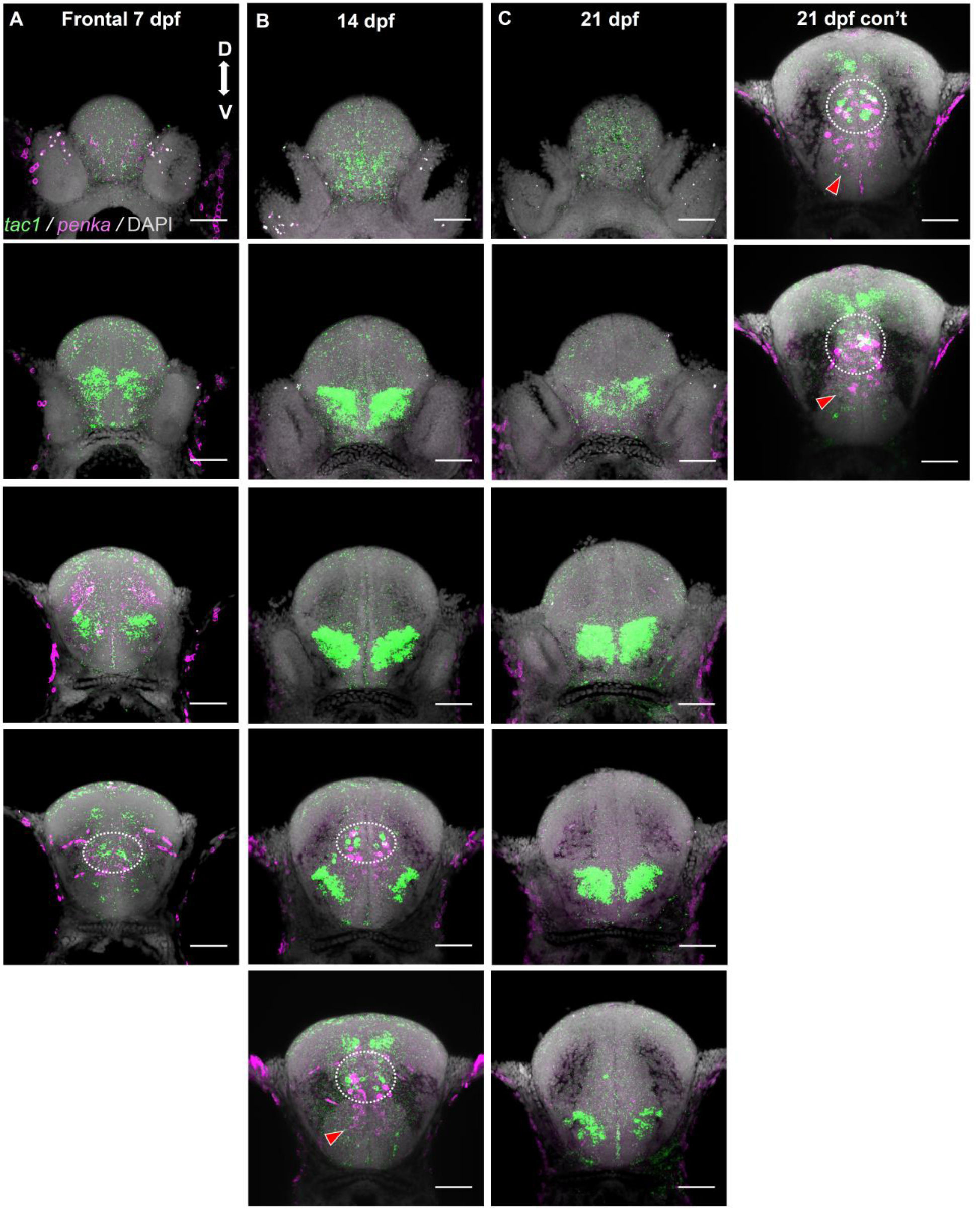
Frontal view of double fluorescent *in situs* for *tac1* and *penka* during larval developement. Confocal z-projections (20 µm) from the forebrain from 7 dpf **(B),** 14 dpf **(C**), and 21 dpf **(D)** larvae. Sections are organized from anterior to posterior, with the most anterior sections in the top row (**A, B, C, D**). White hashed circle indicates area of *tac1* and *penka* comingling. D, dorsal; V, ventral. Scale bar = 50μm.

**Graph 3.4.1:**
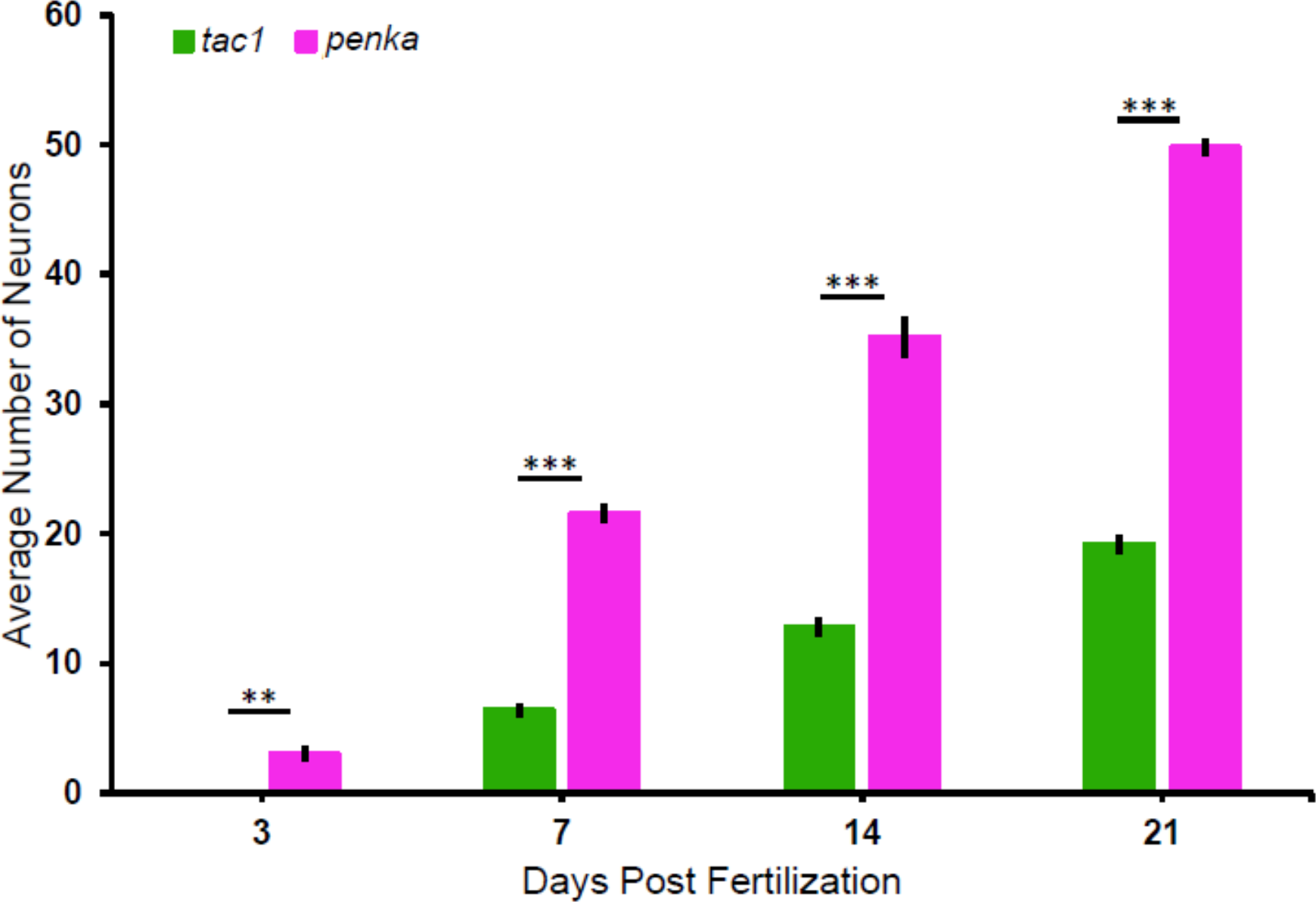
Comparison of *tac1* and *penka* neurons at 3, 7, 14, and 21 dpf. Subpallial *penka* neuronal population is significantly larger than the *tac1* neuronal population. Two-tailed unpaired t-tests were performed, with statistically significant results were reported as p < 0.01 with Bonferroni correction. *** p < 0.0001. ** p < 0.001. Data are represented as mean ± SEM.

**Graph 3.4.2:**
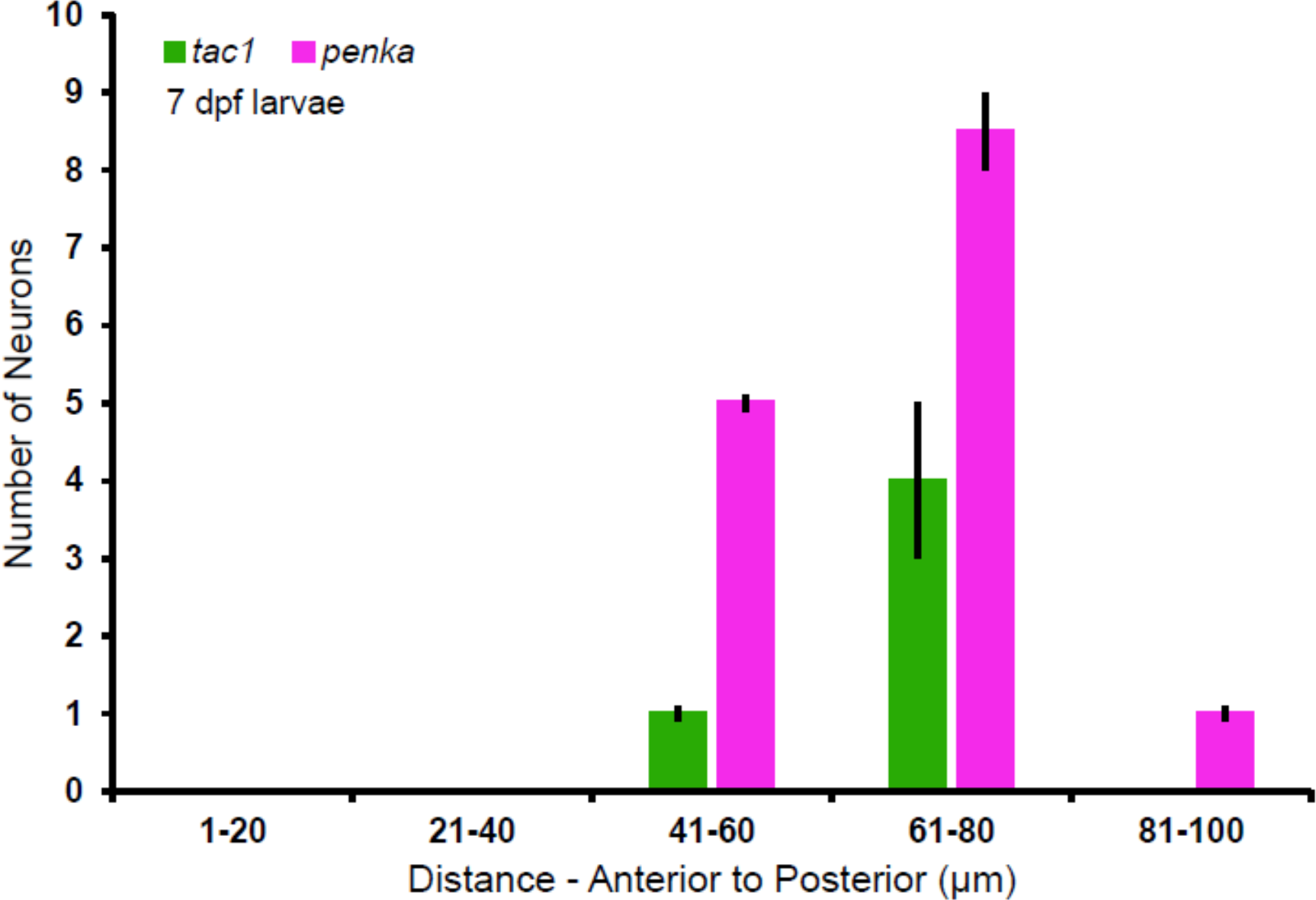
Average number of *tac1* and *penka* neurons from 7 dpf double *in situ* experiments. *tac1* and *penka* cell counts from the subpallium (n = 2). Section 81-100 μm are post-commissural with *penka* neurons likely to belong to preoptic regions. These neurons were not included in the total cell counts of Graphs 3.3.1 or 3.4.1. Data are represented as mean ± SEM.

**Graph 3.4.3:**
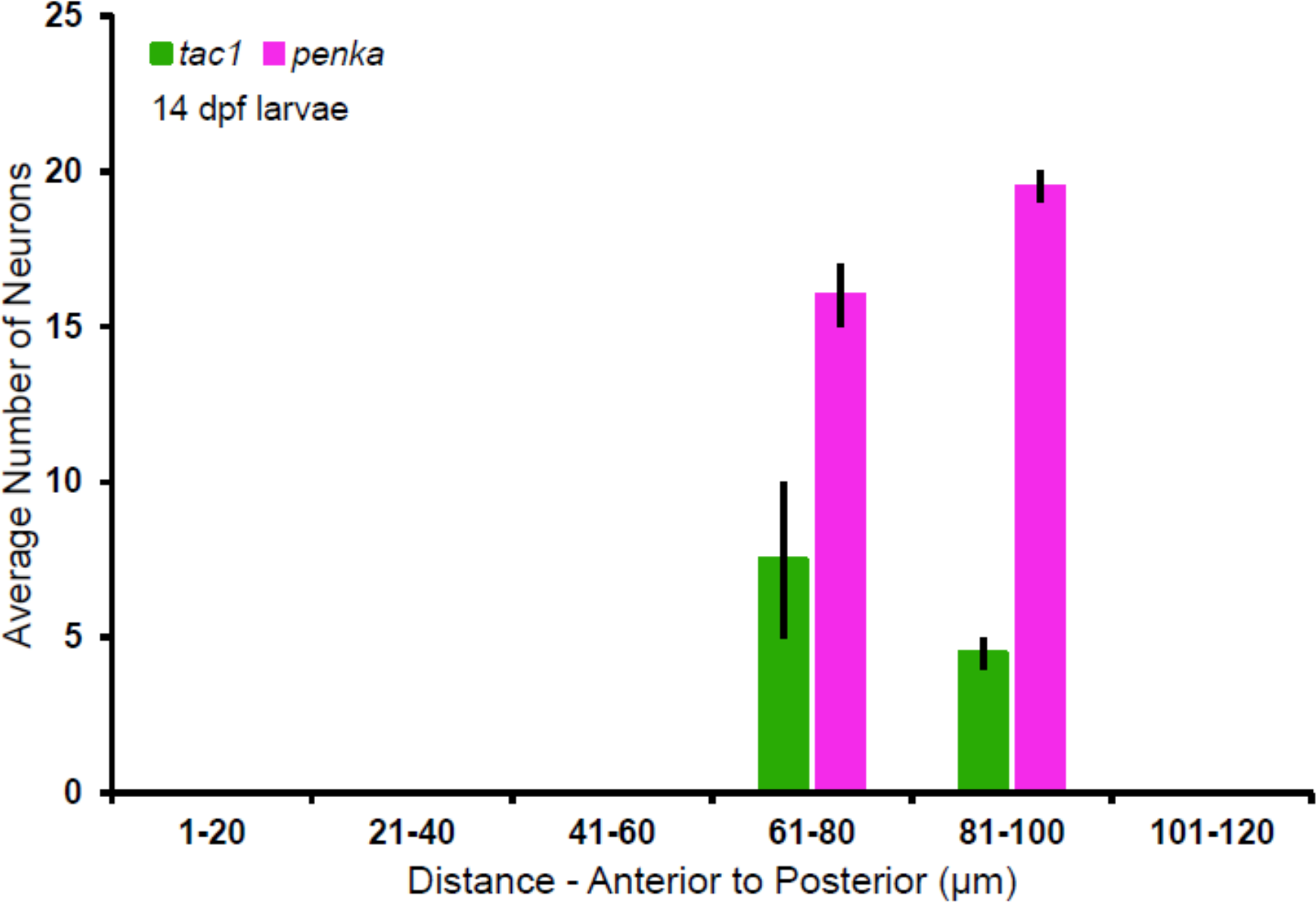
Average number of *tac1* and *penka* neurons from 14 dpf double *in situ* experiments. *tac1* and *penka* cell counts from the subpallium (n = 2). Data are represented as mean ± SEM.

**Graph 3.4.4:**
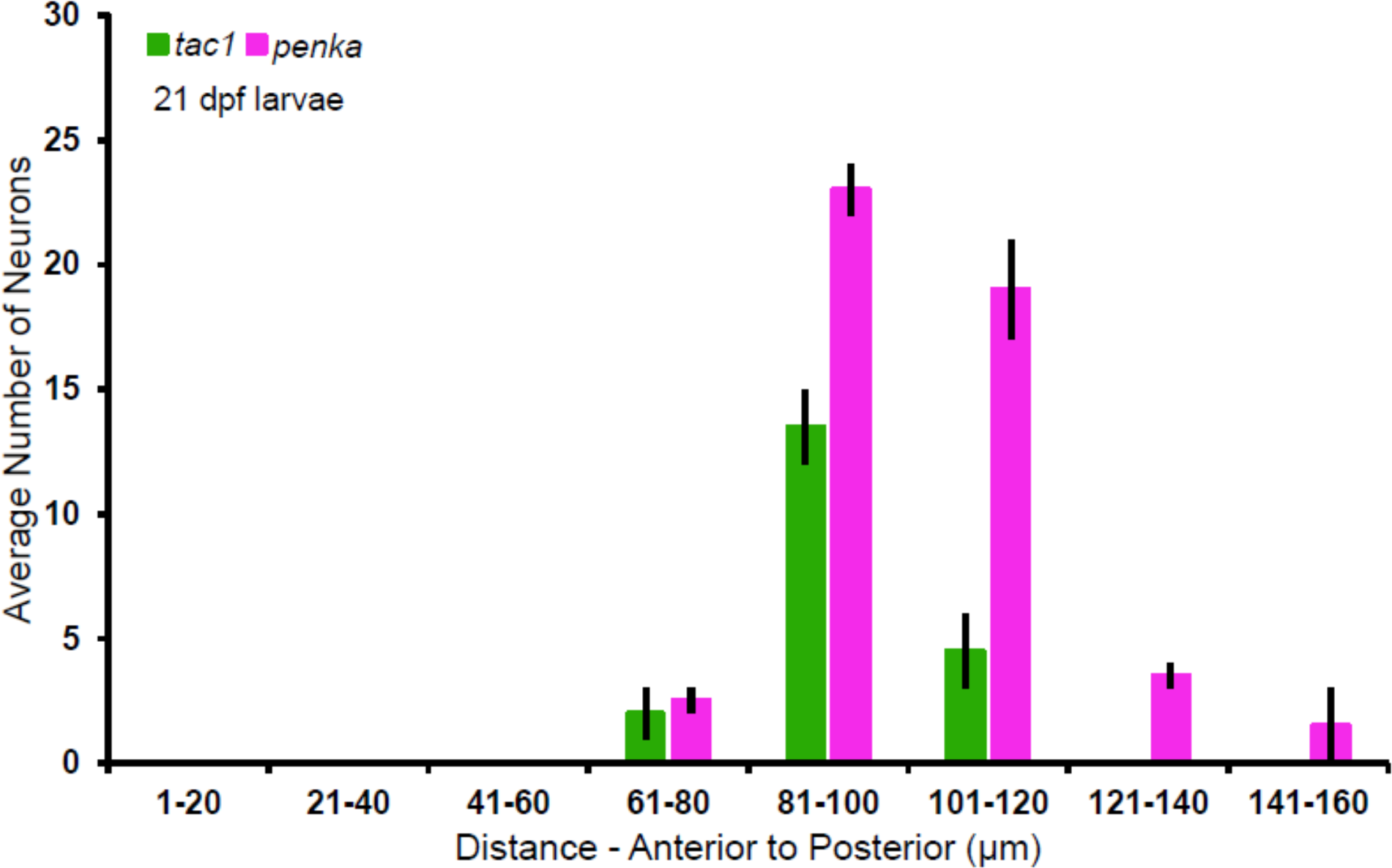
Average number of *tac1* and *penka* neurons from 21 dpf double *in situ* experiments. *tac1* and *penka* cell counts from the subpallium (n = 2). Section 141-160 μm are post-commissural with *penka* neurons likely to belong to preoptic regions. These neurons were not included in the total cell counts of Graphs 3.3.1 or 3.4.1. Data are represented as mean ± SEM.

**Table 3.4.2:** Number of *tac1* and *penka* neurons in precommissural subpallium at 7 dpf, 14 dpf, and 21 dpf from double *in situ* experiments.

|  | *tac1* neurons | *penka* neurons<br>within *tac1*<br>boundary | Total *penka* neurons |
| --- | --- | --- | --- |
| 7 dpf | 6 | 4 | 14 |
|  | 4 | 4 | 15 |
| Average | 5 | 4 | 14.5 |
| 14 dpf | 15 | 12 | 37 |
|  | 9 | 8 | 34 |
| Average | 12 | 10 | 25.5 |
| 21 dpf | 18 | 18 | 48 |
|  | 21 | 22 | 48 |
| Average | 19.5 | 20 | 48 |

**Table 3.4.3:** Number of *tac1* and *penka* neurons in 20 μm sections starting from the anterior most point of the forebrain until the anterior commissure from double *in situ* experiments.

|  |  | 1-20µm | 21-40µm | 41-60µm | 61-80µm | 81-100µm | 101-120µm | 121-140µm | 141-160µm |
| --- | --- | --- | --- | --- | --- | --- | --- | --- | --- |
| **7**<br>**dpf** | *tac1* | 0 | 0 | 1 | 5 | 0 |  |  |  |
|  |  | 0 | 0 | 1 | 3 | 0 |  |  |  |
|  | *penka* | 0 | 0 | 5 | 9 | 1 |  |  |  |
|  |  | 0 | 0 | 5 | 8 | 1 |  |  |  |
| **14**<br>**dpf** | *tac1* | 0 | 0 | 0 | 10 | 5 | 0 |  |  |
|  |  | 0 | 0 | 0 | 5 | 4 | 0 |  |  |
|  | *penka* | 0 | 0 | 0 | 17 | 20 | 0 |  |  |
|  |  | 0 | 0 | 0 | 15 | 19 | 0 |  |  |
| **21**<br>**dpf** | *tac1* | 0 | 0 | 0 | 1 | 15 | 6 | 0 | 0 |
|  |  | 0 | 0 | 0 | 3 | 12 | 3 | 0 | 0 |
|  | *penka* | 0 | 0 | 0 | 3 | 24 | 17 | 4 | 3 |
|  |  | 0 | 0 | 0 | 2 | 22 | 21 | 3 | 0 |

### 3.5 Development of Dopaminergic Neurons Within the Subpallium

Dopamine (DA) is an important modulator of BG signaling. Tyrosine hydroxylase (TH) is often used to identify dopaminergic neurons in the CNS because TH is the rate limiting enzyme in catecholamine synthesis (Molinoff & Axelrod, 1971). TH staining could indicate the presence of dopaminergic or noradrenergic neurons, however, extensive work on these cell types in zebrafish has differentiated these populations (Filippi *et al.,* 2010). We performed TH staining to determine if the precommissural subpallial area is innervated by dopaminergic fibers. We stained whole mount samples with a TH antibody at 3, 7, 14, and 21 dpf. At 3 dpf, there was strong staining of the olfactory bulbs (Figure 3.5.1, A; Figure 3.5.2, column A) with ∼7 neurons lining the pallial/ subpallial boundary (n = 3, Figure 3.5.2, column A, panels 4-5). The olfactory bulbs continued to have strong expression at 7 dpf (Figure 3.5.1, B; Figure 3.5.2, column B, panels 1-2). A TH population of ∼16 neurons (n = 3) was seen extending caudally towards the anterior commissure, remaining at the pallial/ subpallial boundary (Figure 3.5.2, column B, panels 4-7). Stained fibres within the anterior commissure were observed (Figure 3.5.1, B). Similarly at 14 dpf, there was strong staining of the olfactory bulbs (Figure 3.5.1, C) as well as ∼20 TH neurons lining the pallial/ subpallial boundary (n = 3, Figure 3.5.2, column C, panels 4-8). The population of TH positive neurons lining the pallial/ subpallial boundary increased to ∼32 neurons (n = 3) in 21 dpf larvae (Figure 3.5.2, column D, panels 5-9). There was a dense network of stained fibres coursing through the anterior commissure and a scattered population was seen dorsal to the anterior commissure in the dorsal subpallium extending into the thalamus (Figure 3.5.1, D). The dense network of TH stained projections present in this area suggests that the precommissural subpallium receives dopaminergic innervation.

**Figure 3.5.1:**
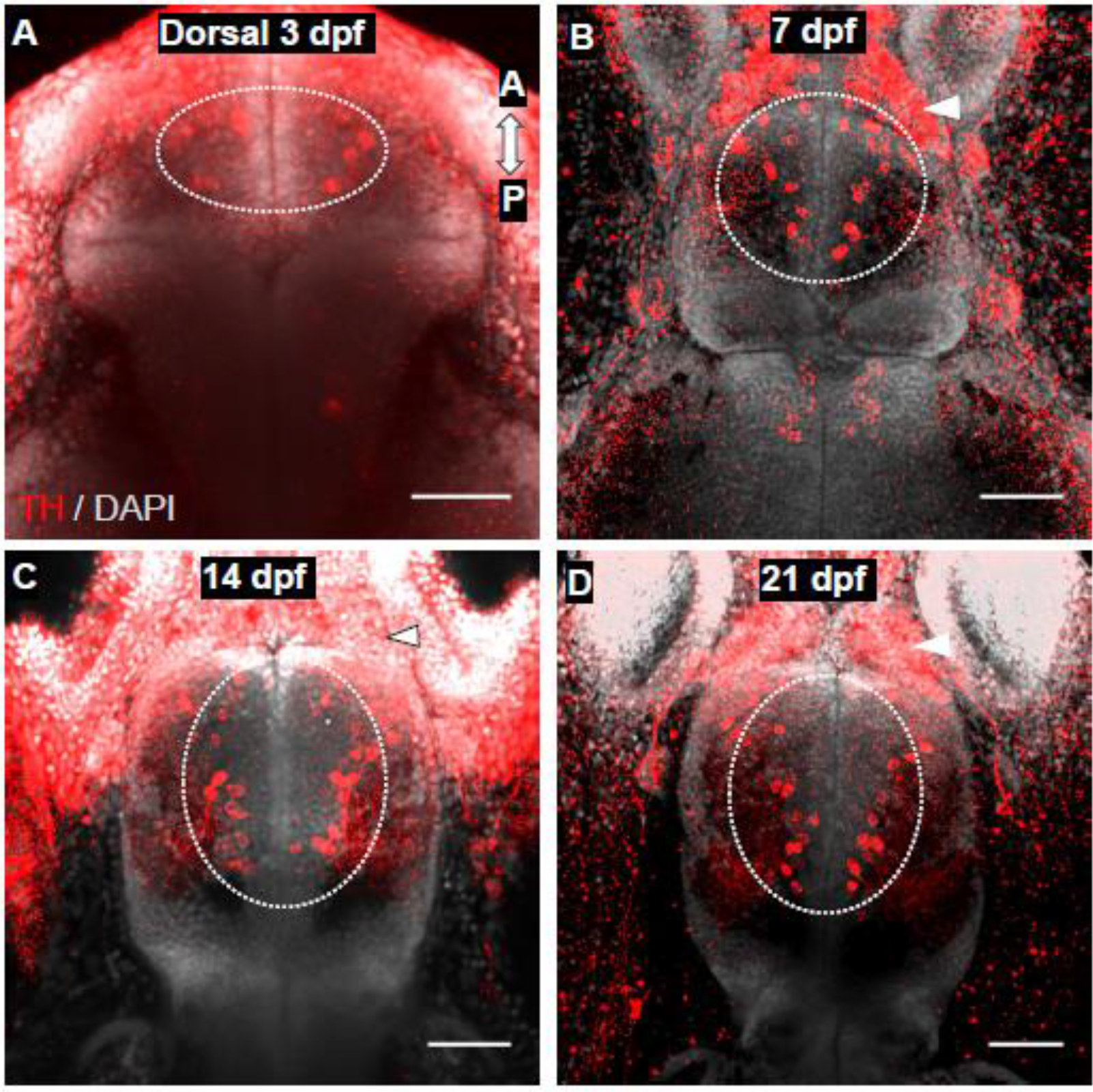
Dorsal view of tyrosine hydroxylase (TH) staining in the forebrain of developing larvae. Confocal z-projections of the forebrain of 3 dpf **(A)**, 7 dpf **(B),** 14 dpf **(C**), and 21 dpf **(D)** larvae. TH staining is present in the olfactory bulb, anterior commissure, and precommissural subpallium. White arrowheads indicate staining in the olfactory bulb. White hashed circle indicates precommissural TH neurons. A, anterior; P, posterior. Scale bar = 50 μm.

**Figure 3.5.2:**
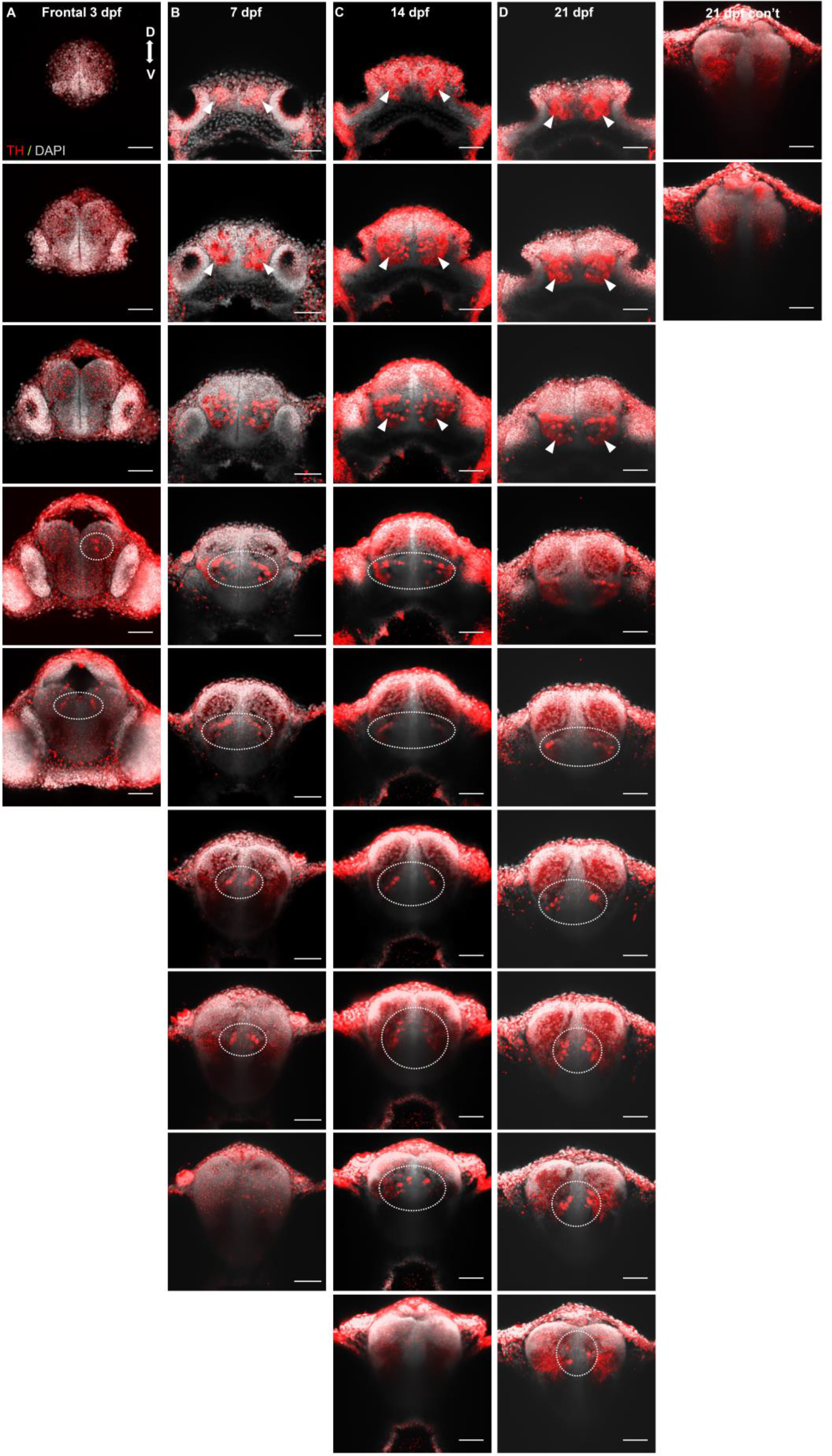
Frontal view of tyrosine hydroxylase (TH) neurons in the subpallium of 3, 7, 14, and 21 dpf larvae. Confocal z-projections (20 µm) of the forebrain of 3 dpf **(A)**, 7 dpf **(B),** 14 dpf **(C**), and 21 dpf **(D)** larvae. Sections are organized from anterior to posterior, with the most anterior sections in the top row (**A, B, C, D**). White arrowheads indicate staining in the olfactory bulb. White hashed circle indicates precommissural TH neurons. D, dorsal; V, ventral. Scale bar = 50 μm.

### 3.6 Dopamine Receptor Gal4 Driver Lines

To begin to functionally characterizing these circuits, we have made efforts to make transgenic Gal4 knock-ins at the dopamine type 1 receptor (D1) locus and dopamine type 2 receptor (D2) locus. These receptors are hypothesized to target the direct and indirect pathways, respectively, as observed in other vertebrates (Surmeir *et al.,* 2007). The Zebrafish CORE facility at SickKids made a presumed insertion of Gal4 at the D1 locus using CRISPR/Cas9 genomic integration (Figure 2.4.2). Subsequently, we engineered a presumed insertion of Gal4 at the D2 locus (Figure 2.4.3). PCR mapping of the potential D1 and D2 insertions has been problematic and we are now pursuing whole genome sequencing to confirm the location of the Gal4 insertions. Both lines have enticing Gal4 expression patterns, with extensive labeling within the subpallium (Figure 3.6.1, A). Potential D1 neurons were present in the olfactory bulb and precommisural subpallial area in 7 dpf larvae. There was extensive expression within fibres traversing through the pallium, anterior commissure, and lateral forebrain bundles. We observed a similar expression pattern within the subpallium for the potential D2 line (Figure 3.6.1, B).

**Figure 3.6.1:**
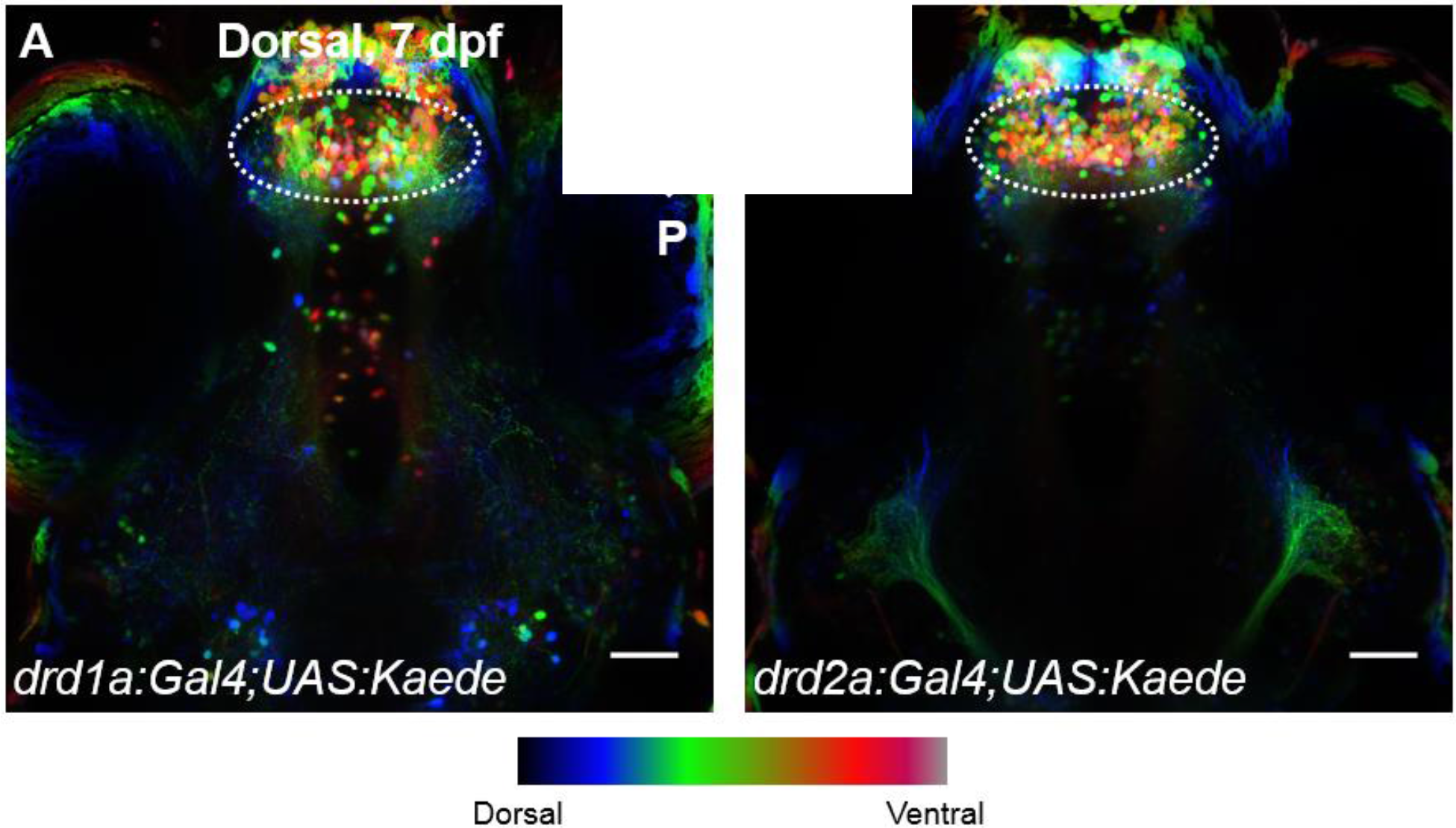
Dorsal view of unverified *drd1a* and *drd2a* transgenic lines. Confocal z-projections of the forebrain of our *drd1a:Gal4;UAS:Kaede* (**A**) and *drd2a:Gal4;UAS:Kaede* **(B)** lines in 7 dpf larvae colour coded for depth. White hashed circle indicates subpallial neurons. A, anterior; P, posterior. Scale bar = 50 μm.

## Discussion

Our study sought to identify circuitry homologous to the mammalian striatum over multiple developmental timepoints in larval zebrafish. We have validated the presence of known striatal markers, including: the presence of GABAergic neurons, substance P expression, enkephalin expression, dopaminergic innervation as well as *tac1* and *penka* gene expression. Additionally, our *tac1* and *penka* time series analyses revealed that both populations expand by more than two-fold from 7 dpf to 21 dpf. These results fell in line with our hypotheses and this research strongly supports the presence of rapidly expanding striatal circuitry in larval zebrafish.

### 4.1 Subpallial Characterization and Location

We used the expression of the homeobox gene, *dlx2a*, to determine the subpallial boundary in 3 dpf larvae. As previously reported, zebrafish *dlx2a* transcripts are expressed in the dorsal and ventral subpallium (Figure 3.1.1; Figure 3.1.2). *dlx2a* expression continues above and caudal to the anterior commissure with the dorsal subpallium. There are laterally positioned *dlx2a* neurons that emerge from the subpallial boundary into the telencephalic region, with some invading the pallium. Previous work shows that the subpallium is divided into dorsal and ventral regions, with the dorsal subpallium giving rise to telencephalic BG components (the most dorsal of the dorsal subpallium being homologous to the caudatoputamen, and the ventral area of the dorsal subpallium being homologous to the pallidum) and the ventral region giving rise to the septum (Mione *et al.,* 2001; Ganz *et al.,* 2012). Our markers show that this putative striatal population is close to the midline of the brain. This differs from most vertebrates and validates findings from published studies stating that the molecular striatal profile in zebrafish is near the dorsal subpallial midline (Ganz *et al.,* 2012).

*dlx2a* plays an important role in the differentiation and migration of most GABAergic neurons within the telencephalon (Anderson *et al.,* 1997). The core neuronal makeup of the mammalian striatum is GABAergic neurons. Within the subpallium of zebrafish at 3, 7, 14, and 21 dpf, we have shown an abundant GABAergic population within the dorsal subpallium (Figure 3.1.4), which is a similar situation in the rodent LGE (Katrova *et al.,* 2000). After 3 dpf, we notice GABAergic neurons extending dorsolaterally at the anterior commissure (Figure 3.1.4, column A, panel 5) into the pallium. In rodents, GABAergic neurons are born in the embryonic ventral subpallium (specifically the MGE), where a large fraction migrate tangentially into the pallium and become interneurons. There are also GABAergic neurons born within the LGE that migrate into the pallium in a late phase of tangential migration. Pallial GABAergic expression was also observed in zebrafish (Mueller *et al.,* 2006). Early expression of GABAergic neurons within the subpallium versus the pallium is a pattern that occurs in amphibians (Brox *et al.,* 2003) and lamprey (Melendez-Ferro *et al.,* 2002) as well, showing that these developmental pathways are conserved amongst vertebrates. Since GABAergic neurons are the basic unit of the BG, this provides further evidence to the conserved nature of the BG and its circuitry amongst vertebrates.

### 4.2 Characterization of the Putative Direct Pathway in Larval Zebrafish

The direct and indirect pathways are key features of BG circuitry and anatomy. Mammalian MSNs of each pathway differ in neuropeptide content and connectivity, with direct pathway neurons containing SP and dynorphin and projecting directly to BG output structures, and indirect pathway neurons containing ENK and projecting indirectly to downstream structures (Gerfen *et al.,* 1992; Charron *et al.,* 1995; Barami *et al.,* 2001). Previous studies alluded to the precommissural dorsal region of the dorsal subpallium in teleosts as being homologous to the mammalian striatum based on histochemical data (Mueller *et al.,* 2008; Ganz *et al.,* 2012). We conducted molecular profiling experiments using precursors to SP and ENK, *tac1* and *penka* respectively, to determine the location of the putative teleost striatum.

Within the subpallium, there are two main areas of *tac1* expression: dense bilateral expression at the anterior most end of the forebrain and a population in the dorsal region of the precommissural dorsal subpallium. We determined the anterior most population to be associated with the olfactory bulbs based on our *tac1 in situs* and SP staining (Figure 3.2.5, D, E). This finding agrees with previously published results from adult zebrafish (Ogawa *et al.,* 2012). *tac1,* and subsequently SP, expression in the olfactory bulb has been reported in various mammals (Sanides-Kohlrausche & Wahle, 1991; Olpe *et al.,* 1987) and is found to depress neuronal activity within the bulbs by releasing GABA. We saw this *tac1* population as early as 3 dpf, whereas the dorsal subpallial population was not seen until 7 dpf. The olfactory system in zebrafish has been shown to be functional before its development is complete and at 2-3 dpf odor responses in the olfactory bulbs can be detected (Miyasaka *et al.,* 2013). At that stage, larvae spend most of their time resting at the bottom of the tank, moving reflexively without directional stability (Muller & van Leeuwen, 2004).

The *tac1* dorsal subpallial population increases as larvae reach ages when they start to swim and perform complex behaviours such as prey capture. At late larval timepoints, sparse *tac1* neurons were present in more ventral areas of the subpallium (Figure 3.4.3, column A, panel 4; column B, panel 5; column C, panel 7). As aforementioned, the ventral portion of the subpallium is believed to give rise to the septum, an evolutionarily conserved part of the limbic system (Mione *et al*., 2001; Ganz *et al.,* 2012). SP expression was restricted to fibres within the olfactory bulbs, anterior commissure, lateral forebrain bundles, pallium, and habenula. Similar to the *tac1* staining, the ventral portion of the dorsal subpallium had very little SP staining.

### 4.3 Characterization of the Putative Indirect Pathway in Larval Zebrafish

*penka* neurons are located along the midline extending along the dorsoventral axis of the subpallium. At 3 dpf, the majority of *penka* neurons are found in lateral telencephalic areas and the preoptic region caudal to the anterior commissure. By 7 dpf onwards, the *penka* population increases at each timepoint, forming a column in the precommissural subpallial area. This expression pattern is similar to primates, where ENK is expressed in more ventral areas of the BG, with relatively strong expression along the dorsoventral axis (Martin & Cork, 2014). The number of *penka* neurons in the dorsal subpallium, especially within the ventral region, increases at each timepoint observed. The ventral portion of the dorsal subpallium is hypothesized to be homologous to the GP, characterized by *nkx2.1* and *lhx6* expression (Manoli & Driever, 2014). While we do not have distinct boundaries for this region, we believe that a portion of the *penka* population is part of the pallidal-region based on their location.

This significant increase in neurons at each timepoint suggests the existence of a pallidal-like region that is potentially online after 7 dpf. In mammals, the globus pallidus contains two different segments: the internal segment (GPi) which receives efferents from SP containing striatal neurons of the direct pathway, and the external segment (GPe) which receives efferents from ENK striatal neurons of the indirect pathway (Charron *et al.,* 1995; Barami *et al.,* 2001). This level of compartmentalization within the globus pallidus is not seen in lower order vertebrates (Marin *et al.,* 1997). It is also hypothesized that since aspects of the indirect pathway (like the GPe and STN) have only been conclusively identified in advanced vertebrates, it may be a more recent evolutionary addition to BG circuitry (Stephenson-Jones *et al.,* 2011). Our data falls in line with that hypothesis, suggesting an all-encompassing GP structure that receives a multitude of different signals from striatal neurons.

### 4.4 Comparison and Development of the Putative Direct and Indirect Pathways in Zebrafish

There is an overlap between *tac1* and *penka* populations in the dorsal region of the precommissural dorsal subpallium that was shown through double FISH labelling at 7, 14, and 21 dpf. Due to a portion of *penka* neurons potentially belonging to pallidal homologs, we used the dorsal *tac1* population to represent the striatal sector. In this area, the number of *tac1* and *penka* neurons appear similar (Table 3.4.2) and comprise a total striatal-like neuronal population of 9 neurons in 7 dpf larvae, 22 neurons in 14 dpf larvae, and 40 neurons in 21 dpf larvae. Using *tac1* neurons as a location marker for the putative striatal region provides a potential boundary between the striatal and pallidal regions in teleosts. The *tac1* and *penka* neuronal populations were intermingled, but distinct, much like what is found in amphibians, reptiles, and birds (Reiner & Northcutt, 1992; Trabucchi *et al.,* 1999; Papalopulu & Kintner, 1993; Tombol *et al.,* 1988). In mammals, however, there is compartmentalization which creates distinct zones called the striosomes within the underlying matrix. These compartments differ in connectivity, neuropeptide content, and function; with the striosomes containing SP and sending efferents to the SNc and the matrix containing ENK and preferentially projecting to the GPe, and SNr (Graybiel, 1990; Charron *et al.,* 1995; Barami *et al.,* 2001).

The lack of compartmentalization in teleosts suggests the existence of a simpler and potentially more streamlined BG circuit, converging sensorimotor, associative, and limbic information onto a striatal network of a countable number of neurons. This highlights the power of using zebrafish for circuit pathway analyses. The population increase seen across timepoints could be attributed to the overall expansion of the brain as the larvae develops. Another reason for this increase could be that as the fish ages, differences in environmental experiences could trigger the expression of *tac1* and *penka* within the precommissural subpallium, as is the case in mammals (Tajima & Fukuda, 2013; Fu & Beckstead, 1992; Wan *et al.,* 1992). The overall increase of both populations coincides with the development of complex behavior and suggests that the complexity of BG circuits within the fish increases as the behavioural repertoire becomes more advanced.

### 4.5 Dopamine Innervation

Dopaminergic innervation of BG nuclei is conserved across evolution, but the location and development of dopaminergic cell groups that project to these nuclei differ between anamniotes and amniotes (Reiner, 2016). TH is the rate-limiting enzyme for catecholamine biosynthesis and serves as a marker for dopaminergic and noradrenergic neurons (Molinoff & Axelrod, 1971). At all timepoints, we observed dense staining of dopaminergic fibres within the olfactory bulb, pallium, anterior commissure, and lateral forebrain bundles. Dopaminergic neurons were found as early as 3 dpf along the pallial-subpallial boundary, where rostral neurons were positioned laterally and caudal neurons closer to the anterior commissure were positioned medially. These local dopaminergic neurons have been reported previously (Tay *et al.,* 2011) and are thought to contribute to dopaminergic signaling between diencephalic structures and BG nuclei. Previous studies have shown a dopaminergic population in the rostral part of the posterior tuberculum that projects to the dorsal subpallium. This population is thought to be similar to A9 neurons found in the SNc of mammals. (Rink & Wullimann, 2001; Tay *et al.,* 2011; Matsui & Sugie, 2017). Our staining suggests that dopaminergic fibres and neurons are present within the subpallium; however, it is unclear whether dopaminergic neurons within the subpallium or posterior tuberculum are modulating BG circuitry. Future studies using neuronal antero- and retrograde tracing of these dopaminergic populations will be required to make this determination.

### 4.6 Development of Transgenic and Computational Tools for the Study of Putative Striatal Circuitry

D1 and D2 receptors are important for modulating the BG circuit. D1 receptors are found on GABAergic neurons of the direct pathway and usually initiate movement, whereas D2 receptors are found on GABAergic neurons of the indirect pathway and usually inhibit movement (Albin *et al.,* 1989; Gerfen *et al.,* 1992). Our unverified transgenic lines show similar expression patterns with the potential for overlapping neuronal populations being labelled. If this turns out to be the case, it would indicate that a portion of subpallial neurons express both D1 and D2 receptors. In mammals there are cases where this occurs (∼1.9% in the dorsal striatum, and ∼22% in the ventral striatum; Gagnon *et al.,* 2017), however the functional significance of coexpression within the overall BG circuitry remains controversial.

In silico techniques provide a means through which we can combine expression data across histological techniques onto a single brain space. The ANTs registration pipeline we developed should allow us to combine our different datasets and make direct comparisons of labeled neuronal populations. In addition, refinement of our brain registration pipeline will allow for future studies that correlate brain activity data (calcium imaging) with the location of direct and indirect pathways markers identified in this thesis. Lastly, the use of brain registration will allow us to add our anatomical data to zebrafish brain atlas databases (Randlett *et al.,* 2015; Tabor *et al.,* 2019).

### 4.7 Limitations and Future Directions

One of the major limitations to this study is the lack of anatomical landmarks/delineations between our processed samples, coupled with the changes in brain morphology that occurs during *in situ* hybridization processing compared to immunostaining. While the subpallial area in zebrafish is relatively clear, the boundaries between the dorsal and ventral subdivisions are not. This could be amended in a number of ways: a) performing *dlx2a* and *nkx2.1* staining for all of the timepoints as a reference for subpallial subdivisions, b) performing *dlx2a* and *nkx2.1* staining for all the time points alongside *tac1* and *penka* staining through triple in situ hybridization techniques, or c) *in silico* brain registration techniques merge expression patterns onto a reference brain. Due to the incompatibility of our RFP antibody with *in situ* processing, we were unable to show *tac1* and *penka* staining in the *gad1b:|R|-GFP* line. We plan on overcoming this with brain registration, which will be able to combine immunostaining with *in situ* staining in the same brain space. We also plan to create additional Gal 4 driver lines labelling the *tac1* and *penka* populations. This would allow for functional experiments including calcium imaging, laser ablations, and optogenetic studies that could determine the role these neurons play in action selection and motor control.

Further characterization of zebrafish striatal cell types would involve examining expression patterns in the adult brain. The transparency of the zebrafish brain decreases with age, but techniques such as CLARITY (Chung & Deisseroth, 2013) and iDisco (Renier *et al.,* 2014) clear tissue allowing for fluorescent signals to be detected in thick tissues. Combining these techniques with the approaches developed in this thesis would allow for a detailed anatomical study of the adult zebrafish striatum.

### 4.8 Conclusion

We used a variety of histological approaches to validate the presence of: 1) GABAergic neurons, 2) SP neurons, 3) ENK neurons, and 4) dopaminergic innervation within the larval zebrafish precommissural dorsal subpallium. We also performed a detailed longitudinal examination of striatal marker expression at stages when zebrafish behaviour develops from purely reflexive outputs to complex social interactions. The characterization of an anatomical region in larval zebrafish that is homologous to the mammalian striatum provides a foundation upon which behavioural and functional analyses of the entire striatum can be built upon. With the addition of Gal4 driver lines that label the zebrafish striatum, it would be technically feasible to optically monitor and manipulate activity in all neurons within the striatum. This would be a feat not possible in any other model organism. Due to the unique systems neuroscience approaches offered in zebrafish, such studies should provide new insights into evolutionarily conserved mechanisms for the selection and execution of behavioural motor programs.

